# Recurrent emergence and transmission of a SARS-CoV-2 spike deletion H69/V70

**DOI:** 10.1101/2020.12.14.422555

**Authors:** Steven A Kemp, Bo Meng, Isabella ATM Ferriera, Rawlings Datir, William T Harvey, Guido Papa, Spyros Lytras, Dami A Collier, Ahmed Mohamed, Giulia Gallo, Nazia Thakur, The COVID-19 Genomics UK (COG-UK) Consortium, Alessandro M Carabelli, Julia C Kenyon, Andrew M Lever, Anna De Marco, Christian Saliba, Katja Culap, Elisabetta Cameroni, Luca Piccoli, Davide Corti, Leo C James, Dalan Bailey, David L Robertson, Ravindra K. Gupta

## Abstract

SARS-CoV-2 amino acid replacements in the receptor binding domain (RBD) occur relatively frequently and some have a consequence for immune recognition. Here we report recurrent emergence and significant onward transmission of a six-nucleotide out of frame deletion in the S gene, which results in loss of two amino acids: H69 and V70. We report that in human infections ΔH69/V70 often co-occurs with the receptor binding motif amino acid replacements N501Y, N439K and Y453F, and in the latter two cases has followed the RBD mutation. One of the ΔH69/V70+ N501Y lineages, now known as B.1.1.7, has undergone rapid expansion and includes eight S gene mutations: RBD (N501Y and A570D), S1 (ΔH69/V70 and Δ144) and S2 (P681H, T716I, S982A and D1118H). *In vitro*, we show that ΔH69/V70 does not reduce serum neutralisation across multiple convalescent sera. However, ΔH69/V70 increases infectivity and is associated with increased incorporation of cleaved spike into virions. ΔH69/V70 is able to compensate for small infectivity defects induced by RBD mutations N501Y, N439K and Y453F. In addition, replacement of H69 and V70 residues in the B.1.1.7 spike reduces its infectivity and spike mediated cell-cell fusion. Based on our data ΔH69/V70 likely acts as a permissive mutation that allows acquisition of otherwise deleterious immune escape mutations. Enhanced surveillance for the ΔH69/V70 deletion with and without RBD mutations should be considered as a global priority not only as a marker for the B.1.1.7 variant, but potentially also for other emerging variants of concern. Vaccines designed to target the deleted spike protein could mitigate against its emergence as increased selective forces from immunity and vaccines increase globally.

**Highlights:**

- ΔH69/V70 is present in at least 28 SARS-CoV-2 lineages
- ΔH69/V70 does not confer escape from convalescent sera
- ΔH69/V70 increases spike infectivity and compensates for RBD mutations
- ΔH69/V70 is associated with greater spike cleavage
- B.1.1.7 requires ΔH69/V70 for optimal spike cleavage and infectivity

## Background

SARS-CoV-2’s spike surface glycoprotein engagement of human angiotensin-converting enzyme (hACE2) is essential for virus entry and infection(Zhou et al., 2020), and the receptor is found in respiratory and gastrointestinal tracts(Sungnak et al., 2020). Despite this critical interaction and the constraints it imposes, it appears the RBD, and particularly the receptor binding motif (RBM), is relatively tolerant to mutations(Starr et al., 2020b; Thomson et al., 2020), raising the real possibility of virus escape from vaccine-induced immunity and monoclonal antibody treatments. Spike mutants exhibiting reduced susceptibility to monoclonal antibodies have been identified in *in vitro* screens(Greaney et al., 2020; Starr et al., 2020a), and some of these mutations have been found in clinical isolates(Choi et al., 2020). Due to the high levels of susceptibility of the human population to this virus, the acute nature of infections and limited use of vaccines to date there has been limited selection pressure placed SARS-CoV-2(MacLean et al., 2020); as a consequence few mutations that could alter antigenicity have increased significantly in frequency.

The unprecedented scale of whole genome SARS-CoV-2 sequencing has enabled identification and epidemiological analysis of transmission and surveillance, particularly in the United Kingdom. As of February 16^th,^ there were 544,778 SARS-CoV-2 sequences available in the GISAID database (https://www.gisaid.org/). However, geographic coverage is very uneven with two-fifths of all sequences being provided by the United Kingdom. Indeed, as more countries continue to present data from samples collected up to six months prior, this may result in novel variants with altered biological or antigenic properties evolving and not being detected until they have already become established at high frequency.

Studying SARS-CoV-2 chronic infections can give insight into virus evolution that would require many chains of acute transmission to generate. This is because the majority of infections arise as a result of early transmission during pre or asymptomatic phases prior to peak adaptive responses, and virus adaptation not observed as the virus is usually cleared by the immune response(He et al., 2020; Mlcochova et al., 2020c). We recently documented *de novo* emergence of antibody evasion mutations mediated by S gene mutations in an individual treated with convalescent plasma (CP)(Kemp et al., 2021). Dramatic changes in the prevalence of Spike variants ΔH69/V70 (an out of frame six-nucleotide deletion) and D796H variant followed repeated use of CP, while *in vitro* the mutated ΔH69/V70+D796H variant displayed reduced susceptibility to CP using a pseudotyped lentivirus assay. D796H alone resulted in more than five-fold loss of infectivity and the ΔH69/V70 partially rescued this defect. In addition, a chronically infected immune suppressed individual was recently reported in Russia with emergence of Y453F, along with ΔH69/V70(Bazykin et al., 2021). Deletions in other parts of the N-Terminal Domain (NTD) have been reported to arise in chronic infections(Choi et al., 2020) and to reduce sensitivity to NTD-specific neutralising antibodies(McCallum et al., 2021; McCarthy et al., 2021).

Here we analyse the variation in the global SARS-CoV-2 data and find ΔH69/V70 occurs independently often emerging after a significant RBD amino acid replacement such as N501Y, Y453F and N439K, that are known to either facilitate neutralising antibody escape or modulate affinity for the human ACE2 receptor. We show that ΔH69/V70 and other common NTD deletions occur at loop structures in RNA where polymerase activity is often compromised. Although structural modelling indicates the ΔH69/V70 is in an exposed loop that contracts post deletion, potentially altering an antigenic site, we report that the ΔH69/V70 does not confer reduced susceptibility to convalescent sera. Functionally, we reveal that ΔH69/V70 does increase Spike infectivity and compensates for an infectivity defect resulting from RBD replacements N501Y, N439K and Y453F. The infectivity increase is associated with higher levels of cleaved S in pseudotyped virions. Finally, the deletion is required for optimal infectivity of the 501Y.V1 (B.1.1.7) spike protein and repair of the two amino acids leads to reduced S incorporation into virions. These data support a role for ΔH69/V70 in promoting virus infectivity to balance deleterious escape mutations and should be mitigated against.

## Results

### Multiple occurences and transmission of Spike ΔH69/V70 with and without S mutations

The deletion ΔH69/V70 is present in over 87,000 sequences worldwide (71,500 from the UK), and has seen global expansion, particularly across much of Europe, Africa and Asia (Figure 1). ΔH69/V70 is observed in at least 28 different global lineages based on PANGO classification, though not all represent multiple independent acquisitions (Figure 1A, Supplementary Table 1). While variants with deletions in this region of Spike are observed in GISAID^13^, the earliest unambiguous sequences that include the ΔH69/V70 were detected on a D614 background in January 2020 (USA and Thailand). The earliest ΔH69/V70 detected on a D614G background was in Sweden in April 2020. The prevalence of ΔH69/V70 has since increased since August 2020 (Figure 1B). Further analysis of sequences revealed, firstly, that single deletions of either 69 or 70 were uncommon and, secondly, some lineages of ΔH69/V70 alone were present, as well as ΔH69/V70 in the context of other mutations in Spike, specifically those in the RBM (Figure 1).

**Figure 1.**
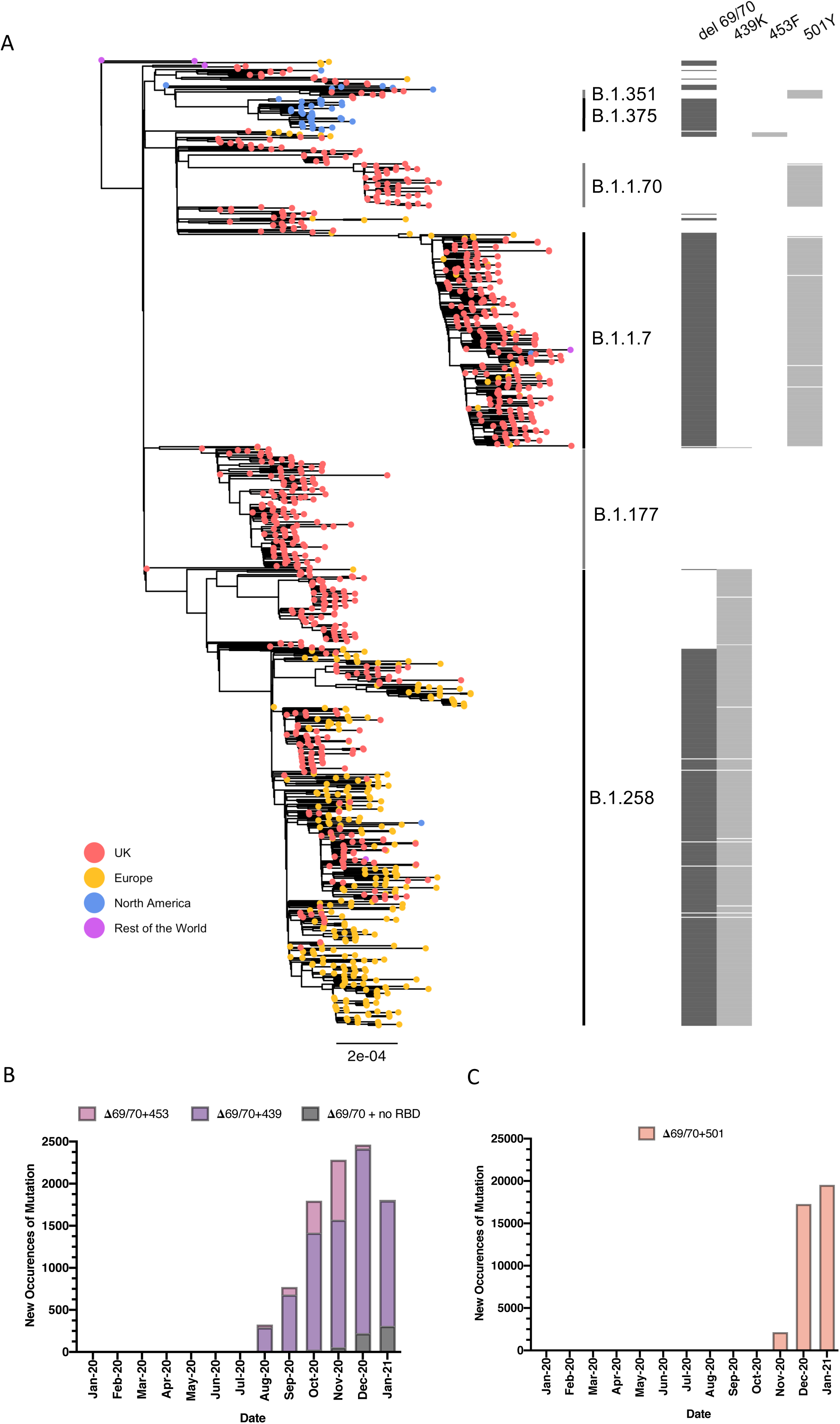
**A. Global phylogeny of SARS-CoV-2 whole genome sequences highlighting those with specific mutations in Spike:** ΔH69/V70, N439K, Y453F and N501Y. Tree is subsampled and tips are coloured by geographic region (see key). Grey bars on the right show the presence or absence of the deletion ΔH69/V70 and amino acid variants N439K, Y453F, and N501Y. Lineages from Rambaut et al. 2020 are shown. **New occurrences of SARS-CoV-2 sequences with the ΔH69/V70 deletion by month for B.** ΔH69/V70 with or without N439K/ Y453F and **C.** ΔH69/V70 with or without N501Y. Indicated frequencies by month of the ΔH69/V70 deletion are from the GISAID database (accessed 18^th^ Feb 2021) by reporting country and sampling date.

We next examined in greater detail the SARS-CoV-2 lineages where S gene mutations in the RBD were identified at high frequency and where ΔH69/V70 co-occurs. For example, N439K, an amino acid replacement reported to be defining variants increasing in numbers in Europe and other regions(Thomson et al., 2020) (Figure 1, Supplementary Figure 1) now mostly co-occurs with ΔH69/V70. N439K appears to have reduced susceptibility to some convalescent sera(Hoffmann et al., 2021) as well as monoclonals targeting the RBD, whilst retaining affinity for ACE2 *in vitro*(Thomson et al., 2020). The first lineage possessing N439K (and not ΔH69/V70), B.1.141 is now extinct(Thomson et al., 2020). A second lineage with N439K, B.1.258, later emerged and subsequently acquired ΔH69/V70 leading to the initial rapid increase in the frequency of viruses possessing this deletion, spreading into (Brejová et al., 2021).

The second significant cluster with ΔH69/V70 and RBD mutants involves Y453F, another Spike RBD mutation that increases binding affinity to ACE2(Starr et al., 2020b) and has been found to be associated with mink-human infections(Munnink et al., 2020). In one SARS-CoV-2 mink-human sub-lineage, termed ‘Cluster 5’, Y453F and ΔH69/V70 occurred with F486L, N501T and M1229I(Larsen et al., 2021) and was shown to have reduced susceptibility to sera from recovered COVID-19 patients (https://files.ssi.dk/Mink-cluster-5-short-report_AFO2). Y453F has also been described as an escape mutation for mAb REGN10933(Baum et al., 2020; Hoffmann et al., 2021). The ΔH69/V70 was first detected in the Y453F background on August 24^th^ 2020 and thus far appears limited to Danish sequences (Figure 1, Supplementary Figure 2), although an independent acquisition was recently reported within an individual, along with ΔH69/V70 in an immune compromised individual with chronic infection(Bazykin et al., 2021).

A third lineage containing the same ΔH69/V70 deletion has arisen with another RBD mutation N501Y along with multiple other spike mutations (Figure 1, Supplementary Figure 3). These UK sequences were subsequently named as belonging a new lineage (B.1.1.7), termed a variant of concern, VOC 202012/01, by Public Health England as it is associated with faster rates of spread (Volz et al., 2021) (Supplementary Figure 3). In addition to RBD N501Y + NTD ΔH69/V70 this new variant had five further S mutations across the NTD (A570D), S2 (P681H, T716I, S982A and D1118H), and NTD 144(Rambaut A., 2020). The variant has now been identified in 94 countries. This lineage has a relatively long branch due to 23 mutations (Supplementary figure 3). The available sequences did not enable us to determine whether the B.1.1.7 mutations N501Y + ΔH69/V70 arose as a result of a N501Y virus acquiring ΔH69/V70 or vice versa, though a sequence was identified with N501Y, A570D, ΔH69/V70 and D1118H, indicating that as predicted N501Y + ΔH69/V70 are proximal mutational events potentially in a long term shedding patient(Kemp et al., 2021). N501Y does escape some RBM targeting antibodies as well as binding ACE2 with higher affinity(Collier et al., 2021a). Sequences with N501Y alone were isolated both in the UK, Brazil, and USA in April 2020, as well as in South Africa.

### ΔH69/V70 and other deletions arise at the terminal end of loop RNA structures

Polymerase processivity can be affected by physical factors in the template caused by RNA structural and sequence motifs and these can facilitate dissociation events. Stable helix loop motifs are associated with pausing/dissociation events in reverse transcriptase(Harrison et al., 1998). Since all nucleic acid polymerases have a common ancestor with homologous dNTP binding motifs and similar global structures(Delarue et al., 1990; Ollis et al., 1985; Sousa et al., 1993) it is probable that all RNA polymerases use similar mechanisms for transcript termination(Reeder and Lang, 1994). Analysis of three deletions in the S protein coding sequence observed across new multiply mutated variants demonstrated that each occurred in the terminal loop of a helix loop motif (Supplementary Figure 4). Although many regions of the structure have a wider structural ensemble of structures that the RNA can adopt, these helix loops were ones in which the ensemble is constrained into a very limited structural range, suggesting that these deletions occur in the more stable stem-loops. A recent in-cell biochemical analysis of SARS-CoV2 RNA structure showed nucleotide reactivity consistent with our model within these stem-loops (Huston et al., 2021). These analyses provide a rationale for preferential emergence of ΔH69/V70 and other deletions such as the well described NTD-antibody escape deletion Y144(Chi et al., 2020; McCarthy et al., 2020, 2021) (in B.1.1.7 and the recently reported B.1.525) at the terminal loops of helical loop motifs.

### ΔH69/V70 does not confer reduced susceptibility to convalescent sera

We hypothesised that ΔH69/V70 is an antibody escape mechanism. We first examined the protein structural context of ΔH69/V70 for clues regarding alterations in epitopes (Figure 2A, B). In the absence of experimentally derived structural data for ΔH69/V70, the protein structure of the NTD possessing the double deletion was modelled in silico. The ΔH69/V70 deletion was predicted to alter the conformation of a protruding loop comprising residues 69 to 76, pulling it in towards the NTD (Figure 2B). In the post-deletion structural model, the positions of the alpha carbons of residues either side of the deleted residues, Ile68 and Ser71, were each predicted to occupy positions 2.9Å from the positions of His69 and Val70 in the pre-deletion structure. Concurrently, the positions of Ser71, Gly72, Thr73, Asn74 and Gly75 are predicted to have changed by 6.5Å, 6.7Å, 6.0Å, 6.2Å and 8Å, respectively, with the overall effect of these residues moving inwards, resulting in a less dramatically protruding loop.

**Figure 2.**
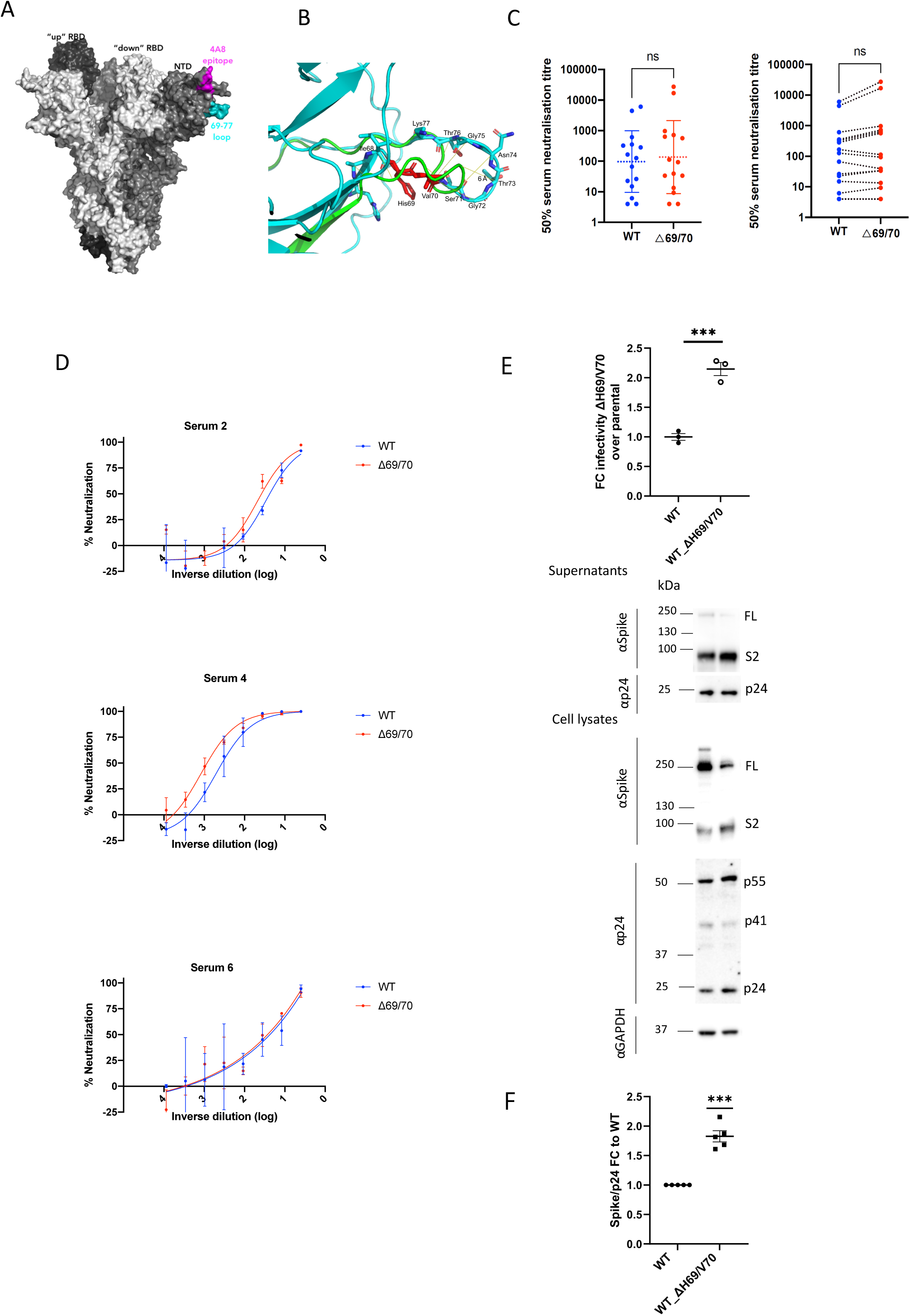
Spike ΔH69/V70 does not reduce sensitivity to neutralising serum antibodies but does increase infectivity. **A.** Surface representation of spike homotrimer in open conformation (PDB: 7C2L) with each monomer shown in different shades of grey. On the monomer shown positioned to the right, the exposed loop consisting of residues 69-77 is shown in cyan and the neutralising antibody (4A8) binding NTD epitope in magenta. **B.** Prediction of conformational change in the spike N-terminal domain due to deletion of residues His69 and Val70. The pre-deletion structure is shown in cyan, except for residues 69 and 70, which are shown in red. The predicted post-deletion structure is shown in green. Residues 66-77 of the pre-deletion structure are shown in stick representation and coloured by atom (nitrogen in blue, oxygen in coral). Yellow lines connect aligned residues 66-77 of the pre- and post-deletion structures and the distance of 6 Å between aligned alpha carbons of Thr73 in the pre- and post-deletion conformation is labelled. **C.** Neutralisation of spike ΔH69/V70 pseudotyped virus and wild type (D614G background) by convalescent sera from 15 donors. GMT (geometric mean titre) with s.d presented of two independent experiments each with two technical repeats. Wilcoxon matched-pairs signed rank test, ns not significant. **D.** Example neutralisation curves. Indicated is serum log_10_ inverse dilution against % neutralisation. Data points represent means of technical replicates and error bars represent standard deviation. **E.** Single round infection by spike ΔH69/V70 pseudotyped virus. Data from three experiments shown with mean and standard error of mean (SEM). Representative western blot of unspun supernatant and cell lysates probed with antibodies for HIV-1 p24 and SARS-Cov-2 S2.**F.** Quantification of cleaved S2 spike:p24 ratio for wild type virus with ΔH69/V70 deletion versus WT alone across multiple replicate experiments. Mean and SEM are shown Students t-test *** p<0.001.

This predicted change in the surface of Spike could be consistent with antibody evasion. To test this we explored whether ΔH69/V70 conferred reduced susceptibility to neutralising antibodies in sera from fifteen recovered individuals (Figure 2C, D). We performed serial dilutions of sera before mixing with viral particles pseudotyped with Spike proteins with and without ΔH69/V70 (with virus input normalised for infectivity). We plotted infection of target cells as a function of serum dilution across the fifteen serum samples (Figure 2D, Supplementary Figure 5). All but two sera demonstrated clear titratable neutralisation of both wild type and ΔH69/V70 virus. There was no overall change in susceptibility to serum neutralisation for ΔH69/V70 relative to wild type (Figure 2C), but there were a proportion of individuals with increase in neutralisation to ΔH69/V70 with small shift in the titration curves (Figure 2C, D, Supplementary Figure 5). In these cases the ΔH69/V70 was associated with increased susceptibility to sera. These data suggest that ΔH69/V70 does not represent an important antibody escape mechanism.

### ΔH69/V70 spike infectivity correlates with increased cleaved spike in virions

Given the association between ΔH69/V70 and RBD mutations and data from an earlier report in chronic infection^12^, we hypothesised that this deletion might alternatively enhance virus infectivity. In the absence of virus isolates we used a lentiviral pseudotyping approach to test the impact of ΔH69/V70 on virus Spike protein mediated infection. A D614G bearing Spike protein expressing DNA plasmid (wild type, WT) was co-transfected in HEK 293T producer cell lines along with plasmids encoding lentiviral capsid and genome for luciferase. Infectivity was adjusted for input reverse transcriptase activity; we observed a two-fold increase in infectivity of ΔH69/V70 as compared to WT (Figure 2E). Western blotting indicated that for ΔH69/V70 there was a higher amount of cleaved spike in the virion containing cell supernatants (unspun) and in the producer cell lysates. It appeared that there was a corresponding reduction in uncleaved spike (Figure 2E). Densitometric analysis of spike and p24 from western blots in multiple experiments showed almost a two-fold increase in spike:p24 ratio for the ΔH69/V70, indicating increased spike cleavage efficiency and incorporation into virions that could explain the increase in infectivity. In order to explore whether D614G was required for this enhanced cleavage efficiency and infectivity, we generated pseudotyped virus bearing D614 spike with and without the ΔH69/V70 and infected target cells. We observed a similar enhancement of infection and increase in cleaved spike in supernatants as we did for D614G spike pseudotyped viruses (Supplementary Figure 6).

### Spike ΔH69/V70 compensates for reduced spike infectivity due to RBD replacements

We hypothesised that ΔH69/V70 might compensate for potential loss of infectivity due to receptor binding motif mutations Y453F, N510Y and N439K that interact with ACE2 (Figure 3A-C). We generated mutant spike plasmids bearing RBD mutations N501Y, Y453F, or N439K both with and without ΔH69/V70 (Figure 3A-C) and performed infectivity assays in the lentiviral pseudotyping system. RBD mutations reduced infectivity of Spike relative to WT by 2-3 fold (Figure 3D, E). Based on observations of the impact of ΔH69/V70 on S incorporation, we predicted that the mechanism of increased infectivity for ΔH69/V70 in context of RBD mutations might also involve increased incorporation of cleaved S2 spike as in Figure 2E. The analysis on virions from cell supernatants and cell lysates indeed showed increased cleaved S2 spike when ΔH69/V70 was present with RBD mutants (Figure 3E, F).

**Figure 3:**
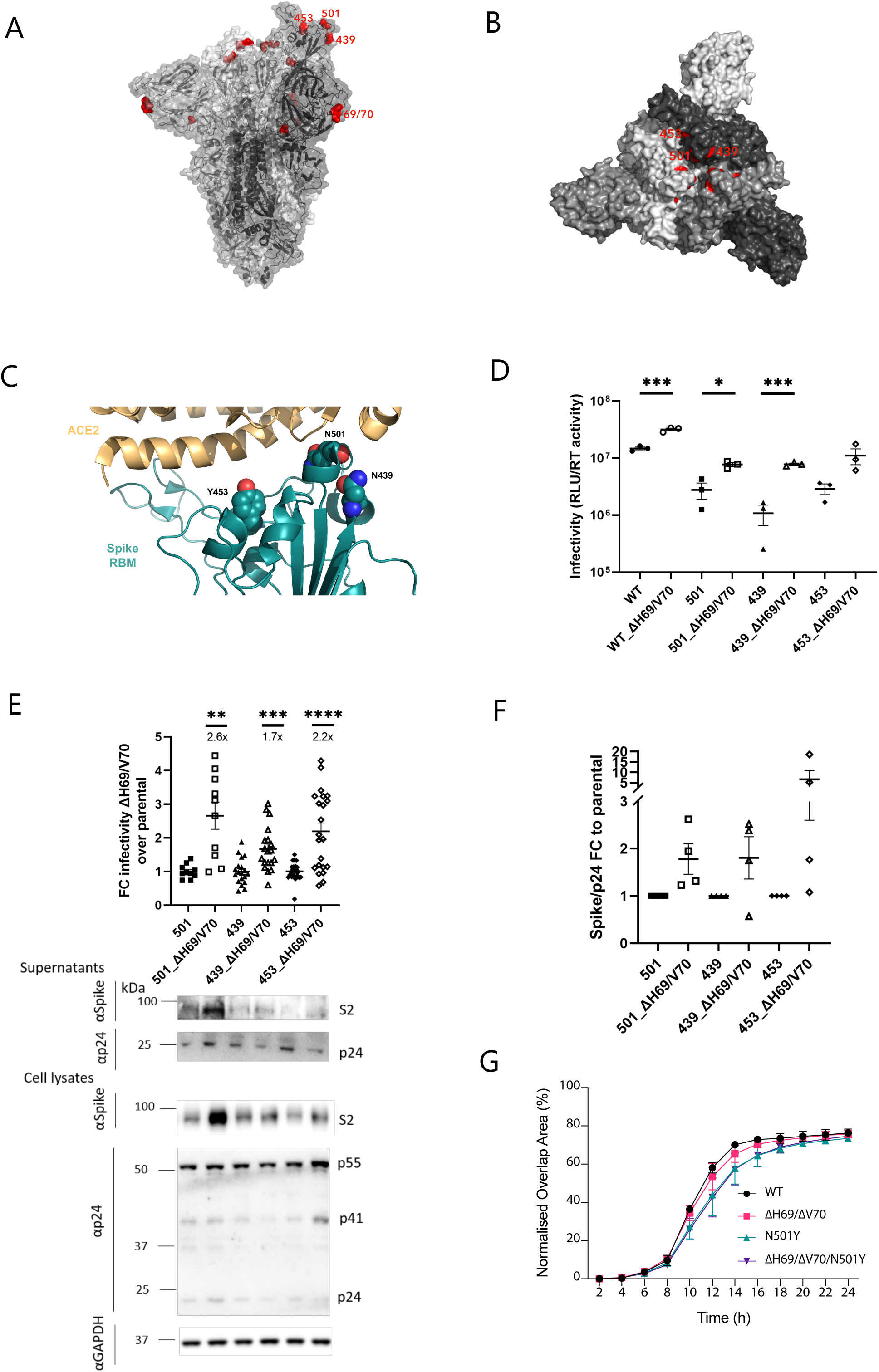
ΔH69/V70 compensates for reduced infectivity of RBD mutations N439K, Y453F and N501Y. **A.** Spike in open conformation with a single erect RBD (PDB: 6ZGG) in trimer axis vertical view with the location of ΔH69/V70 in the N-terminal domain and RBD mutations highlighted as red spheres and labelled on the monomer with erect RBD**. B.** Surface representation of spike in closed conformation (PDB: 6ZGE) viewed in a ‘top-down’ view along the trimer axis. The residues associated with RBD substitutions N439K, Y453F and N501Y are highlighted in red and labelled on a single monomer. **C.** Representation of Spike RBM:ACE2 interface (PDB: 6M0J) with residues N439, Y453 and N501 highlighted as spheres coloured by element. **D-F**. Spike mutant ΔH69/V70 compensates for infectivity defect of Spike RBD mutations and is associated with increased Spike incorporation into virions. **D.** Infectivity of Spike (D614G) ΔH69/V70 deletion in absence and presence of Spike RBD mutations. Single round infection by luciferase expressing lentivirus pseudotyped with SARS-CoV-2 spike protein on HeLa cells transduced with ACE2. **E**. Fold change infectivity over multiple experiments comparing RBD mutants with and without ΔH69/V70, with mean and SEM shown. Representative western blot of unspun supernatant and cell lysates probed with antibodies against HIV-1 p24 and SARS-CoV-2 spike S2. **F**. Quantification of cleaved S2 spike:p24 ratio for wild type virus with ΔH69/V70 deletion versus WT alone across multiple replicate experiments. Mean and SEM are shown G. Cell-cell fusion kinetics for mutant spike proteins. Students t-test *** p<0.001. Data for D and E show infectivity normalized for virus input using reverse transcriptase activity in virus supernatants. RLU – relative light units; U – unit of reverse transcriptase activity (RT). Data are technical replicates and are representative of 2 independent experiments. Student t test * p<0.05, ***p<0.001

In order to explore the mechanism of increased infectivity, we used a cell fusion assay (Figure 3G, Supplementary Figure 7) to monitor kinetics of cell fusion. Previous reports have shown that SARS-CoV-2 Spike protein possesses high fusogenic activity and is able to trigger the formation of large multi-nucleated cells (named syncytia) *in vitro* and *in vivo* (Papa et al., 2020) (Cattin-Ortola at el. 2020), potentially providing an additional and a more rapid route for virus disseminating among neighbour cells. To understand whether the higher infectivity of ΔH69/V70 spike might be related to an increased ability to trigger cell-cell fusion between different cell types, we ectopically overexpressed SARS-CoV-2 spike variants together with the mCherry fluorescent protein in 293T cells and label Vero cells with a green fluorescent dye. Mixing both cell types and measuring the merged green and red fluorescence allows precise quantification of cell-cell fusion kinetics (Supplementary Figure 7). Syncytia formation kinetics appeared slightly lower for N501Y SARS-CoV-2 Spike mutant compared to wild type, ΔH69/V70 and N501Y along with ΔH69/V70 (Figure 3G).

### B.1.1.7 Spike cleavage efficiency, virion incorporation and infectivity is reduced by re-insertion of H69V70

B.1.1.7 naturally contains the ΔH69/V70 deletion (Figure 4A). We predicted that the replacement of H69 and V70 would impair the infectivity of B.1.1.7. To examine this, we compared the infectivity of B.1.1.7 Spike versus B.1.1.7 without the ΔH69/V70 deletion in our pseudotyping system. We observed that infectivity of B.1.1.7 and WT pseudotyped viruses were similar (Figure 4B). As expected, we observed a significant reduction in infectivity for viruses where the H69 and V70 had been re-inserted (Figure 4B, C). When we tested Spike cleavage and incorporation into virions we found that the reduced infectivity of the B.1.1.7 with replaced H69 V70 was associated with reduced cleaved spike S2 protein (Figure 4C, D).

**Figure 4.**
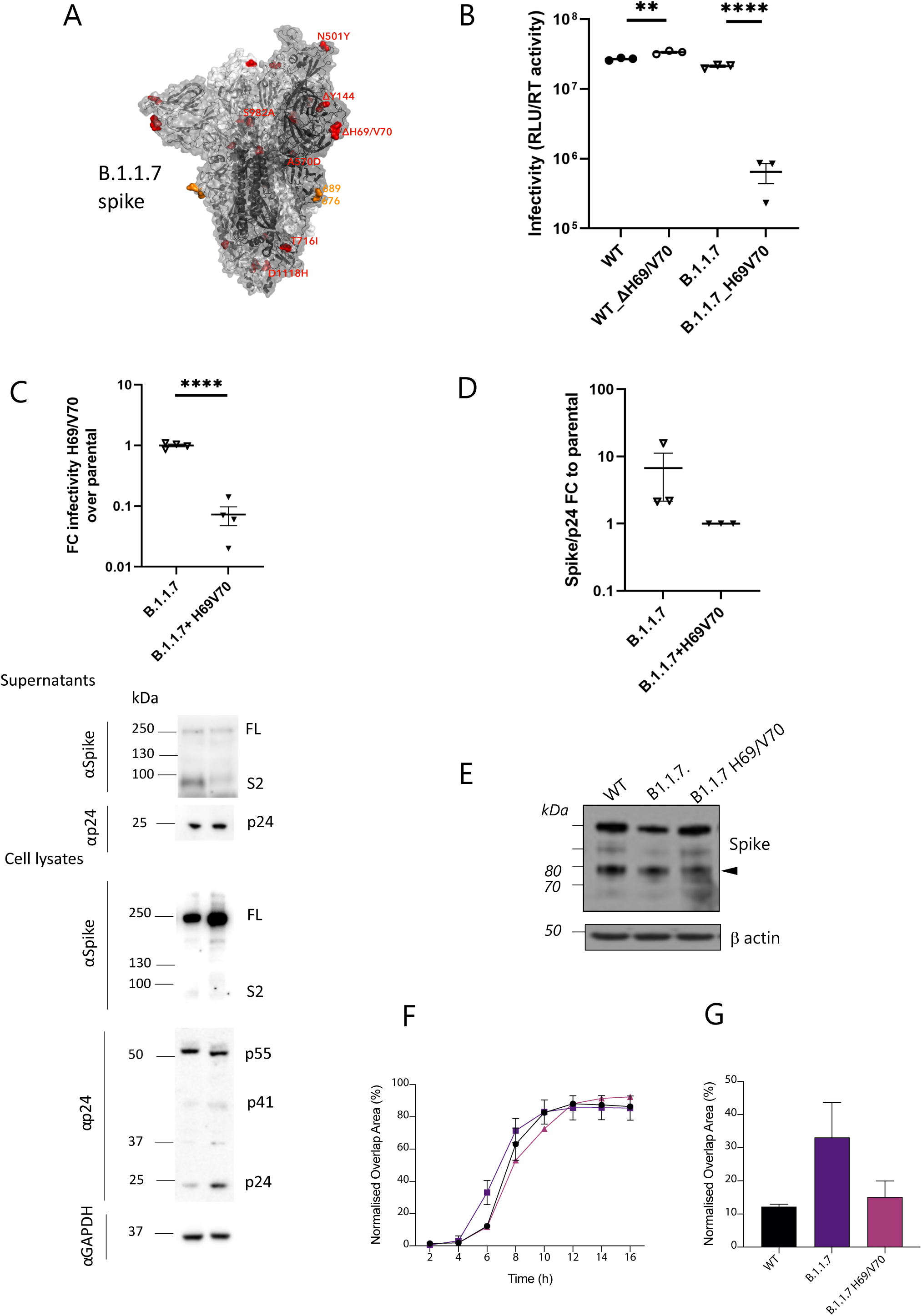
Spike ΔH69/V70 in B.1.1.7 enhances spike infectivity and cell fusogenicity. **A.** Surface representation of spike homotrimer in open conformation with one upright RBD overlaid with ribbon representation (PDB: 6ZGG, Wrobel et al., 2020), with different monomers shown in shades of grey. The deleted residues H69 and V70 and the residues involved in amino acid substitutions (501, 570, 716, 982 and 1118) and the deletion at position 144 are coloured red on each monomer and labelled on the monomer with an upright RBD. The location of an exposed loop containing the furin cleavage site and including residue 681 is absent from the structure, though modelled residues either side of this loop, 676 and 689, are coloured orange. **B.** Infectivity of B.1.1.7 with replacement of H69 and V70 versus B.1.1.7 containing Spike ΔH69/V70 and wild type (D614G) spike. Single round infection by luciferase expressing lentivirus pseudotyped with SARS-CoV-2 Spike protein on HeLa cells transduced with ACE2. **C.** Infectivity of B.1.1.7 without the ΔH69/V70 expressed as fold change compared to B.1.1.7, and representative western blot analysis following transfection of cells with spike and lentiviral plasmids. Supernatant (unspun) loading was normalized for input virus input using reverse transcriptase activity in virus supernatants. **D.** Quantification of Spike:p24 ratio for wild type virus with ΔH69/V70 deletion versus WT alone across replicate experiments. Antibodies against HIV-1 p24 and Spike S2 were used. Data are representative of at least two independent experiments. **E-F. Cell-cell fusion.** Representative western blot of cells transfected with the indicated Spike mutants. The cleaved Spike identifies the S2 subunit and is indicated with the arrowhead. Quantification of cell-cell fusion kinetics showing percentage of green and red overlap area over time, indicative of successful cell-cell fusion. **G.** Data for cell-cell fusion at 6hrs post transfection. Data are representative of two independent experiments

In order to ascertain whether H69V70 represented a target for neutralising antibodies in the context of B.1.1.7, we tested 13 NTD-specific mAbs isolated from 4 individuals that recovered from WT SARS-CoV-2 infection with an *in-vitro* pseudotyped neutralization assay using VeroE6 target cells expressing Transmembrane protease serine 2 (TMPRSS2, **Supplementary Table 1**). The pseudotyped viruses expressed the WT SARS-CoV-2 S, the B.1.1.7 S or the B.1.1.7 S with reversion of ΔH69/V70 deletions (B.1.1.7 ΔH69/V70). We found that 8 out of 13 (62%) showed a marked decrease or complete loss of neutralizing activity to both B.1.1.7 and B.1.1.7 ΔH69/V70 (>30 fold-change reduction), suggesting that in a sizeable fraction of NTD antibodies the ΔH69/V70 deletion is not responsible for their loss of neutralizing activity (Supplementary Figure 8). Only 3 mAbs showed a partial reduction (3- to-8 fold) in B.1.1.7 neutralization that was rescued or improved in one case by reversion of ΔH69/V70 deletions. Neutralization of 2 mAbs (15%) was not affected by both deletion and reversion of ΔH69/V70 in B.1.1.7.

### ΔH69/V70 in sarbecoviruses closely related to SARS-CoV-2

Finally, to investigate the importance of this part of spike beyond SARS-CoV-2, we examined the 69/70 region of spike in a set of other known *Sarbecoviruses* (Figure 5). We observed substantial variability in the region, resulting in frequent indels, with some viruses including SARS-CoV having 6-7 amino acid deletions (Figure 5B). This is indicative of plasticity in this protein region that could allow the *sarbecoviruses* to alter their Spike conformation. The second closest relative to SARS-CoV-2 for this region after RaTG13 is the cluster of 5 CoVs sampled in trafficked pangolins in the Guangxi province(Lam et al., 2020). Inspection of the 69/70 region in these virus sequences raises the interesting observation that one of the five viruses in the cluster, P2V, has amino acids 69H and 70L present, while the other four have a double amino acid deletion (Figure 5B). Given that SARS-CoV-2 and RaTG13 have the homologous HV insertion at these positions, one explanation is that the proximal common ancestor between SARS-CoV-2 and the Guangxi pangolin cluster had the insertion, which was then lost while circulating in the pangolin population, similar to observations with SARS-CoV-2 in humans. Yet, the fact that P2V was cultured in Vero E6 cells prior to sequencing (contrary to the other 4, sequenced directly from the pangolin sample) raises the possibility of this being an independent insertion, favoured as a cell line specific adaptation. Interestingly, the double amino acid indel in the pangolin viruses is in-frame in contrast to SARS-CoV-2 (e.g. lineage B.1.1.7, Figure 5C).

**Figure 5.**
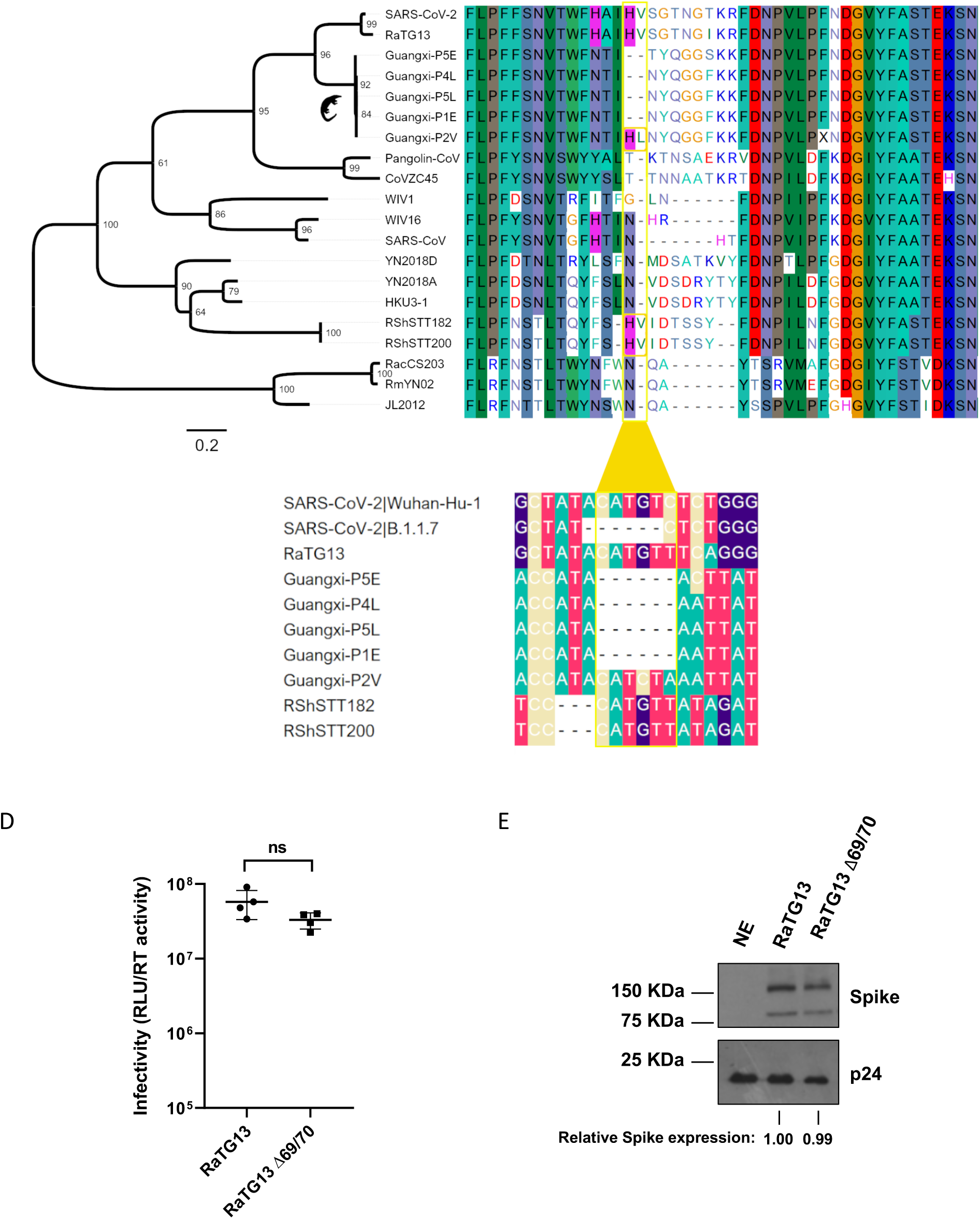
Comparison of the ΔH69/V70 deletion site to other *Sarbecoviruses*. **A**. phylogeny for the Spike peptide region 1-256 **B**. protein sequences from 20 Sarbecoviruses, including SARS-CoV-2 (Wuhan-Hu-1) and SARS-CoV (HSZ-Cc), with distinct genotypes at the Spike region around amino acid positions 69 and 70 (highlighted in yellow box). The 69/70 HL insertion in the P2V sequence from the Guangxi pangolin virus cluster and the HV convergent insertion in the RShSTT182/200 bat virus sequences are highlighted. **C**. The nucleotide alignment between SARS-CoV-2 Wuhan-Hu-1, B.1.1.7, the bat sarbecovirus RaTG13 RShSTT182/200 and the Guangxi pangolin viruses shows the difference between the out-of-frame deletion observed in the former and the in-frame deletion in the latter. **D.** Single round infection by luciferase expressing lentivirus pseudotyped with RaTG13 Spike protein on 293T cells transduced with ACE2. Experiments were performed in biological quadruplicate with the mean and standard deviation plotted. Results are representative of experiments performed two times. Statistical significance was assessed using an unpaired t-test (ns; non-significant, ***; <0.005). **E**. Representative western blot of supernatant from virus producer cells. Spike and HIV pseudotype abundances were assessed using Flag and p24 antibodies, respectively. Relative spike expression was calculated by densitometry using Image J. NE: no envelope.

Furthermore, the two almost identical bat viruses recently sequenced from Cambodia samples – RShSTT182 and RShSTT200 (Hul et al., 2021) possess an H69V70 insertion despite being more distantly related to SARS-CoV-2 for this region of Spike (Figure 5). This independent occurrence of the insertion is suggestive of conditional selective pressures playing a role in recurring gain and loss of these two residues in the sarbecoviruses. To test whether the beneficial effect of ΔH69/V70 is specific to SARS-CoV-2 and not other *Sarbecovirus* Spike backgrounds, we cloned full length S from RaTG13 with a C terminal FLAG tag and generated pseudotyped lentiviruses expressing RaTG13 Spike protein as well as a RaTG13 S H69V70 counterpart. Interestingly, we observed that cleaved and uncleaved S expression levels in unpelleted supernatants containing pseudotyped virus did not differ between WT and the H69V70 RaTG13 spike, and that there was no difference in infectivity. This indicates that the enhancing effect of H69V70 in SARS-CoV-2 may be virus background specific.

## Discussion

We have presented data demonstrating multiple, independent, and circulating lineages of SARS-CoV-2 variants bearing spike ΔH69/V70. This recurring deletion spanning six nucleotides is due to an out of frame deletion of six nucleotides, and occurs in the terminal loop of a helix loop motif within the RNA structure, as do other NTD deletions observed in new variants such as the UK B.1.1.7, South African B.1.351, Brazilian P.1. ΔH69/V70 has frequently followed receptor binding amino acid replacements (N501Y, N439K and Y453F that have been shown to increase binding affinity to hACE2 and reduce binding with monoclonal antibodies), and is specifically found in the B.1.1.7 variant known to have higher transmissibility (Volz et al., 2020)

The ΔH69/V70 deletion was recently shown to emerge following treatment with convalescent plasma^12^. In a second case of multiple mutations in context of immune suppression the ΔH69/V70 deletion occurred with Y453F, though without biological characterisation. We show here experimentally that the ΔH69/V70 deletion is indeed able increase infectivity of viruses bearing RBD mutations N501Y, N439K and Y453F, potentially explaining why the deletion is often observed after these RBD mutation in SARS-CoV-2 global phylogenies. We show that the mechanism of enhanced infectivity across the RBD mutations tested is associated with greater spike cleavage and incorporation of cleaved spike into virions where the ΔH69/V70 deletion is present. Importantly, we were able to recapitulate the ΔH69/V70 phenotype in a spike protein that did not have the D614G mutation, indicating that D614G is not involved in the mechanism of S incorporation enhancement. These observations are supported by ΔH69/V70 being observed in D614 viruses in Jan 2020 both in the US and Thailand.

Critically, we show that whilst B.1.1.7 spike has similar infectivity as wild type D614G spike, there is substantial loss of infectivity when the ΔH69/V70 amino acids are replaced, and this is accompanied by reduced S incorporation into virions. These data point to co-evolution of the observed mutations in spike of B.1.1.7, with a balance of mutations that incur fitness cost with those that aid immune evasion. The findings suggest that the ΔH69/V70 may contribute to the higher viral loads and increased transmissibility of B.1.1.7. Taken together these data support a model whereby ΔH69/V70 can act as a ‘permissive’ mutation that enhances infection, with the potential to enhance the ability of SARS-CoV-2 to tolerate immune escape mutations that would have otherwise significantly reduced viral fitness.

Of note, whilst we showed that RBD mutations reduced infectivity in a pseudotyping system, this reduced infectivity may not translate directly to vivo replication where ACE2 levels on target airway cells are lower. For example the higher affinity of N501Y for ACE2 may be selected for by low ACE2 levels, but incur a fitness defect in higher ACE2 environments. Alternatively, increased ACE2 affinity afforded by 501Y may confer a selective advantage under immune pressure, such as exposure to treatment by convalescent plasma, but require compensatory mutations to transmit optimally among naïve hosts. In the successful lineages B.1.351 and P.1, which also carry 501Y, it is notable that 501Y is accompanied by RBD substitutions K417N and K417T respectively, each of which reduce ACE2 affinity (Starr et al. 2020); potentially acting as alternative compensatory mutations for 501Y.

However, we do believe that the effect of ΔH69/V70 is robust; firstly, our in vitro work involving removal and insertion of the ΔH69/V70 residues significantly impacts both cleaved spike protein levels in virions and infectivity; secondly, the ΔH69/V70 has emerged and transmitted as a single mutation and as a co-mutation in viruses that have previously acquired an RBD mutation such as Y453F, R439K and N501Y, arguing for a role in transmissibility.

The potential for SARS-CoV-2 to evolve and fix mutations is exemplified by D614G, an amino acid replacement in S1 that alters linkages between S1 and S2 subunits on adjacent protomers as well as RBD orientation, infectivity, and transmission(Hou et al., 2020; Korber et al., 2020; Yurkovetskiy et al., 2020). While D614G confers a transmissibility advantage, the resulting increased tendency for the open conformation of spike results in increased susceptibility to neutralisation by RBD-binding monoclonal antibodies (Weissmann et al., 2020). The example of D614G also demonstrates that mechanisms directly impacting important biological processes can be indirect. Similarly, a number of possible mechanistic explanations may underlie ΔH69/V70. For example, the fact that it sits on an exposed surface and is estimated to alter the conformation of a particularly exposed loop might be suggestive of immune interactions and escape, though we have presented data to show that ΔH69/V70 did not reduce sensitivity of spike to neutralising antibodies in serum from a group of recovered individuals. Indeed in some sera, susceptibility to ΔH69/V70 was increased relative to wild type, raising the hypothesis that, similar to D614G, ΔH69/V70 simultaneously increases infectivity and increases susceptibility to neutralising antibodies by a conformational change favouring a more open spike conformation. Consistent with these data, Xie et al recently reported increased susceptibility of N501Y + ΔH69/V70 full length viruses(Xie et al., 2021) in a proportion of post vaccine sera, and our recent work on B.1.1.7 spike showed increased susceptibility of a triple mutant bearing N501Y + A570D + ΔH69/V70 to both vaccine and convalescent sera(Collier et al., 2021b).

Allosteric interactions, as postulated for D614G, could lead to the higher cleavage efficiency and infectivity of spike pseudotyped viruses with ΔH69/V70. It is also possible that the increased infectivity of relates to interactions with receptors other than ACE2 or TMPRSS2, for example L-SIGN/ DC-SIGN. (Soh et al., 2020)Notably, data on increased infectivity conferred by D614G using similar pseudotyped viruses(Yurkovetskiy et al., 2020) was also observed in whole virus and translated to increased viral load and transmission in animal models(Hou et al., 2020). It would be important to investigate whether ΔH69/V70 is associated with higher viral loads or increased transmission, though such epidemiological studies are highly complex and prone to confounding/bias.

The finding of a lineage (B.1.1.7), termed VOC 202012/01, bearing seven S gene mutations across the RBD (N501Y, A570D), S1 (ΔH69/V70 and 144) and S2 (P681H, T716I, S982A and D1118H) in UK requires further experimental characterisation. The detection of a high number of novel mutations suggests this lineage has either been introduced from a geographic region with very poor sampling or viral evolution may have occurred in a single individual in the context of a chronic infection(Kemp et al., 2020). The emergence of variants with higher numbers of mutations so far in the UK and South Africa may herald an era of re-infection and threaten future vaccine efficacy if left unchecked. Perhaps of greatest concern is the emergence of E484K on the background of B.1.1.7. B.1.1.7 possesses a clear transmissibility advantage(Volz et al., 2021) and considerably changes antigenic profile across spike with escaping a range of NTD-targeting neutralising antibodies (McCarthy et al. 2021, McCallum et al. 2021) and E484K which escapes a range of RBD-targeting neutralising antibodies (Baum et al. 2020, Liu et al. 2021, Greaney et al. 2021).

The presence of sequence at site 69/70 appears to be unique to SARS-CoV-2, the closest bat sarbecovirus, RaTG13, recently identified bat viruses in Cambodia RShSTT182 and RShSTT200, and one of the Guangxi pangolin sequences. Although we cannot delineate between the indel being lost in the pangolin host, or gained after culturing the virus, both scenarios are suggestive of functional importance at this protein region. In contrast to SARS-CoV-2, deletion of ΔH69/V70 in the bat *sarbecovirus* RaTG13 did not impact S incorporation or infectivity. This could be explained by the lack of a polybasic cleavage site in these *sarbecoviruses,* virus genetic backgrounds, or cellular factors not recapitulated in human cell lines.

The detection and surveillance of B.1.1.7 has been facilitated in the UK by the phenomenon of SGTF (S gene target failure) due to primers in the Thermofisher SARS-CoV-2 diagnostic qPCR assay used by a significant number of testing facilities. The S gene target (binding in the region of ΔH69/V70) is one of three and therefore a marker for the spread of B.1.1.7 has been tracked by the loss of signal in the S gene target(Volz et al., 2021). However recent reports from the US and central Europe caution against use of SGTF as a sole marker for B.1.1.7 as a significant ΔH69/V70 lineage without other mutations in spike is circulating in the US, and a B.1.258 lineage with N439K with ΔH69/V70, circulating in Slovakia/Czech republic(Brejová et al., 2021; Larsen and Worobey, 2020). Such examples highlight the need for sequencing to accompany novel approaches to diagnostics for variants.

Given the emergence of multiple clusters of variants carrying RBD mutations and the ΔH69/V70 deletion, limitation of transmission takes on a renewed urgency. Continued emphasis on testing/tracing, social distancing and mask wearing are essential, with investment in other novel methods to limit transmission(Mlcochova et al., 2020a). Detection of the deletion and other key mutations by rapid diagnostics should be a research priority as such tests could be used as a proxy for antibody escape mutations to inform surveillance at global scale. Finally, comprehensive vaccination efforts should be accelerated in order to further limit transmission and acquisition of further mutations, and future vaccines could include ΔH69/V70 in order to close this route for virus evolution, assuming that effective neutralising antibodies to this region are generated.

## Limitations

The laboratory virology aspects of this study were conducted with pseudoviruses rather than replication competent viruses. We also carried out experiments in cells overexpressing receptors.

## Acknowledgements

RKG is supported by a Wellcome Trust Senior Fellowship in Clinical Science (WT108082AIA). COG-UK is supported by funding from the Medical Research Council (MRC) part of UK Research & Innovation (UKRI), the National Institute of Health Research (NIHR) and Genome Research Limited, operating as the Wellcome Sanger Institute. This study was supported by the Cambridge NIHRB Biomedical Research Centre. SAK is supported by the Bill and Melinda Gates Foundation via PANGEA grant: OPP1175094. DLR is funded by the MRC (MC UU 1201412). WH is funded by the MRC (MR/R024758/1). We thank Dr James Voss for the kind gift of HeLa cells stably expressing ACE2. SL is funded by Medical Research Council MC_UU_12014/12. This study was also partly funded by Rosetrees Trust. AML is funded by the Cambridge NIHRB Biomedical Research Centre.

## Author contributions

Designed study and experiments: R.K.G, B.M., R.D., I.A.TM.F, D.L.R, L.C.J., D.B., L.P., A.D.M, D.C. Designed and performed structural analysis: W.T.H, A.M.C. Performed experiments: S.A.K, B.M., A.D.M. Interpreted data: L.P., A.D.M, D.C., B.M., R.K.G, D.L.R, D.B, R.D., I.A.TM.F, S.A.K. Carried out pseudovirus neutralization assays: A.D.M. Produced pseudoviruses: C.S. Sequencing and expression of antibodies, mutagenesis for mutant expression plasmids: E.C. and K.C. Analysis of the data and manuscript preparation: L.P., D.C., R.D., I.A.TM.F, R.K.G, B.M., S.A.K.

## Methods

### Phylogenetic Analysis

All available full-genome SARS-CoV-2 sequences were downloaded from the GISAID database (http://gisaid.org)(Shu and McCauley, 2017) on 16^th^ February 2021. Low-quality sequences (>5% N regions) were removed, leaving a dataset of 491,395 sequences with a length of >29,000bp. Sequences were deduplicated and then filtered to find the mutations of interest. All sequences were realigned to the SARS-CoV-2 reference strain MN908947.3, using MAFFT v7.475 with automatic strategy selection and the --keeplength --addfragments options(Katoh and Standley, 2013). Major SARS-CoV-2 clade memberships were assigned to all sequences using the Nextclade server v0.13 (https://clades.nextstrain.org/), Pangolin v2.2.2(Rambaut et al., 2020) (<u>github.com/cov-lineages/pangolin</u>) and a local instance of the PangoLEARN model, dated 17^th^ Feb 2021 21:49 (https://github.com/cov-lineages/pangoLEARN).

Maximum likelihood phylogenetic trees were produced using the above curated dataset using IQ-TREE v2.1.2(Minh et al., 2020). Evolutionary model selection for trees were inferred using ModelFinder(Kalyaanamoorthy et al., 2017) and trees were estimated using the GTR+F+I model with 1000 ultrafast bootstrap replicates(Minh et al., 2013). All trees were visualised with Figtree v.1.4.4 (http://tree.bio.ed.ac.uk/software/figtree/) and ggtree v1.14.6 rooted on the SARS-CoV-2 reference sequence and nodes arranged in descending order. Nodes with bootstraps values of <50 were collapsed using an in-house script.

To reconstruct a phylogeny for the 69/70 spike region of the 20 *Sarbecoviruses* examined in Figure 5, Rdp5(Martin et al., 2015) was used on the codon spike alignment to determine the region between amino acids 1 and 256 as putatively non-recombinant. A tree was reconstructed using the nucleotide alignment of this region under a GTR+_Γ_ substitution model with RAxML-NG(Kozlov et al., 2019). Node support was calculated with 1000 bootstraps. Alignment visualisation was done using BioEdit(Hall et al., 2011).

### RNA secondary structure modelling

2990 nucleotides centred around the spike protein amino acids 69-70 from SARS-CoV2 sequence from an individual^12^ were aligned in CLUSATL-Omega (nucleotides 20277-23265 of the Wuhan isolate MN908947.3) and a consensus structure was generated using RNAalifold(Bernhart et al., 2008)).

### Structural modelling

The structure of the post-deletion NTD (residues 14-306) was modelled using I-TASSER(Roy et al., 2010), a method involving detection of templates from the protein data bank, fragment structure assembly using replica-exchange Monte Carlo simulation and atomic-level refinement of structure using a fragment-guided molecular dynamics simulation. The structural model generated was aligned with the spike structure possessing the pre-deletion conformation of the 69-77 loop (PDB 7C2L(Chi et al., 2020)) using PyMOL (Schrödinger). Figures prepared with PyMOL using PDBs 7C2L, 6M0J(Lan et al., 2020), 6ZGE28 and 6ZGG(Wrobel et al., 2020).

### Cells

HEK 293T CRL-3216, Vero CCL-81 were purchased from ATCC and maintained in Dulbecco’s Modified Eagle Medium (DMEM) supplemented with 10% fetal calf serum (FCS), 100 U/ml penicillin, and 100mg/ml streptomycin. All cells are regularly tested and are mycoplasma free.

### Pseudotype virus preparation

Plasmids encoding the spike protein of SARS-CoV-2 D614 with a C terminal 19 amino acid deletion with D614G, were used as a template to produce variants lacking amino acids at position H69 and V70, as well as mutations N439K, Y453F and N501Y. Mutations were introduced using Quickchange Lightning Site-Directed Mutagenesis kit (Agilent) following the manufacturer’s instructions. B.1.1.7 S expressing plasmid preparation was described previously, but in brief was generated by step wise mutagenesis. Viral vectors were prepared by transfection of 293T cells by using Fugene HD transfection reagent (Promega). 293T cells were transfected with a mixture of 11ul of Fugene HD, 1µg of pCDNAΔ19 spike-HA, 1ug of p8.91 HIV-1 gag-pol expression vector and 1.5µg of pCSFLW (expressing the firefly luciferase reporter gene with the HIV-1 packaging signal). Viral supernatant was collected at 48 and 72h after transfection, filtered through 0.45um filter and stored at −80°C as previously described. Infectivity was measured by luciferase detection in target 293T cells transfected with TMPRSS2 and ACE2.

### SARS-CoV-2 D614 (Wuhan) and RaTG13 mutant plasmids and infectivity

Plasmids encoding the full-length spike protein of SARS-CoV-2 D614 (Wuhan) and RaTG13, in frame with a C – terminal Flag tag(Conceicao et al., 2020), were used as a template to produce variants lacking amino acids at position H69 and V70. The deletion was introduced using Quickchange Lightning Site-Directed Mutagenesis kit (Agilent) following the manufacturer’s instructions. Viruses were purified by ultracentrifugation; 25mL of crude preparation being purified on a 20% sucrose cushion at 2300rpm for 2 hrs at 4°C. After centrifugation, the supernatant was discarded and the viral pellet resuspended in 600 µL DMEM (10% FBS) and stored at −80°C. Infectivity was examined in HEK293 cells transfected with human ACE2, with RLUs normalised to RT activity present in the pseudotyped virus preparation by PERT assay. Western blots were performed on purified virus with anti-HIV1 p24, 1:1,000 (Abcam) or anti-FLAG, 1:2,000 (Sigma) antibodies used following SDS-PAGE and transfer.

### Standardisation of virus input by SYBR Green-based product-enhanced PCR assay (SG-PERT)

The reverse transcriptase activity of virus preparations was determined by qPCR using a SYBR Green-based product-enhanced PCR assay (SG-PERT) as previously described(Vermeire et al., 2012). Briefly, 10-fold dilutions of virus supernatant were lysed in a 1:1 ratio in a 2x lysis solution (made up of 40% glycerol v/v 0.25% Trition X-100 v/v 100mM KCl, RNase inhibitor 0.8 U/ml, TrisHCL 100mM, buffered to pH7.4) for 10 minutes at room temperature.

12µl of each sample lysate was added to thirteen 13µl of a SYBR Green master mix (containing 0.5µM of MS2-RNA Fwd and Rev primers, 3.5pmol/ml of MS2-RNA, and 0.125U/µl of Ribolock RNAse inhibitor and cycled in a QuantStudio. Relative amounts of reverse transcriptase activity were determined as the rate of transcription of bacteriophage MS2 RNA, with absolute RT activity calculated by comparing the relative amounts of RT to an RT standard of known activity.

### Cell-cell fusion assay

Cell fusion assay was carried out as previously described. Briefly, Vero cells and 293T cells were seeded at 80% confluency in a 24 multiwell plate. 293T cells were co-transfected with 1.5 mg of spike expression plasmids in pCDNA3 and 0.5 mg pmCherry-N1 using Fugene 6 and following the manufacturer’s instructions (Promega). Vero cells were treated with CellTracker™ Green CMFDA (5-chloromethylfluorescein diacetate) (Thermo Scientific) for 20 minutes. 293T cells were then detached 5 hours post transfection, mixed together with the green-labelled Vero cells, and plated in a 12 multiwell plate. Cell-cell fusion was measured using an Incucyte and determined as the proportion of merged area to green area over time. Data were then analysed using Incucyte software analysis. Data were normalised to cells transfected only with mCherry protein and mixed with green labelled Vero cells. Graphs were generated using Prism 8 software.

### Transfection

HEK 293T cells were transfected with 1.5 mg of spike expression plasmids in pCDNA3 and lysed 18 hours post transfection. Cells were treated with Benzonase Nuclease (70664 Millipore) and boiled for 5 min. Samples were then run on 4%–12% Bis Tris gels and transferred onto nitrocellulose membranes using an iBlot (Life Technologies).

### Western blots

Cells were lysed with cell lysis buffer (Cell signalling) or were treated with Benzonase Nuclease (70664 Millipore) and boiled for 5 min. Samples were then run on 4%–12% Bis Tris gels and transferred onto nitrocellulose or PVDF membranes using an iBlot or semidry (Life Technologies and Biorad, respectively).

Membranes were blocked for 1 hour in 5% non-fat milk in PBS + 0.1% Tween-20 (PBST) at room temperature with agitation, incubated in primary antibody (anti-SARS-CoV-2 Spike, which detects the S2 subunit of SARS-CoV-2 S (Invitrogen, PA1-41165), anti-GAPDH (proteintech) or anti-p24 (NIBSC)) diluted in 5% non-fat milk in PBST for 2 hours at 4°C with agitation, washed four times in PBST for 5 minutes at room temperature with agitation and incubated in secondary antibody (anti-rabbit or anti-mouse HRP conjugate), anti-bactin HRP (sc-47778) diluted in 5% non-fat milk in PBST for 1 hour with agitation at room temperature. Membranes were washed four times in PBST for 5 minutes at room temperature and imaged directly using a ChemiDoc MP imaging system (Bio-Rad).

### Serum pseudotype neutralisation assay

Spike pseudotype assays have been shown to have similar characteristics as neutralisation testing using fully infectious wild type SARS-CoV-2(Schmidt et al., 2020).Virus neutralisation assays were performed on 293T cell transiently transfected with ACE2 and TMPRSS2 using SARS-CoV-2 spike pseudotyped virus expressing luciferase(Mlcochova et al., 2020b). Pseudotyped virus was incubated with serial dilution of heat inactivated human serum samples or convalescent plasma in duplicate for 1h at 37°C. Virus and cell only controls were also included. Then, freshly trypsinized 293T ACE2/TMPRSS2 expressing cells were added to each well. Following 48h incubation in a 5% CO_2_ environment at 37°C, the luminescence was measured using Steady-Glo Luciferase assay system (Promega).

Ethical approval for use of serum samples. Controls with COVID-19 were enrolled to the NIHR BioResource Centre Cambridge under ethics review board (17/EE/0025).

### Monoclonal antibody neutralisation of B.1.1.7 or B.1.1.7 ΔH69/V70 pseudotyped viruses

Preparation of B.1.1.7 or B.1.1.7 ΔH69/V70 SARS-CoV-2 S glycoprotein-encoding-plasmid used to produce SARS-CoV-2-MLV based on overlap extension PCR. Briefly, a modification of the overlap extension PCR protocol(Forloni et al., 2018) was used to introduce the 9 or 7 mutations of the B.1.1.7 and B.1.1.7 ΔH69/V70 lineages, respectively. In a first step, 9 DNA fragments with overlap sequences were amplified by PCR from a plasmid (phCMV1, Genlantis) encoding the full-length SARS-CoV-2 S gene (BetaCoV/Wuhan-Hu-1/2019, accession number mn908947). The mutations (del-69/70, del-144, N501Y, A570D, D614G, P681H, S982A, T716I and D1118H or K417N, E484K and N501Y) were introduced by amplification with primers with similar Tm. Deletion of the C-terminal 21 amino acids was introduced to increase surface expression of the recombinant S(Case et al., 2020). Next, 3 contiguous overlapping fragments were fused by a first overlap PCR (step 2) using the utmost external primers of each set, resulting in 3 larger fragments with overlapping sequences. A final overlap PCR (step 3) was performed on the 3 large fragments using the utmost external primers to amplify the full-length S gene and the flanking sequences including the restriction sites KpnI and NotI. This fragment was digested and cloned into the expression plasmid phCMV1. For all PCR reactions the Q5 Hot Start High fidelity DNA polymerase was used (New England Biolabs Inc.), according to the manufacturer’s instructions and adapting the elongation time to the size of the amplicon. After each PCR step the amplified regions were separated on agarose gel and purified using Illustra GFX™ PCR DNA and Gel Band Purification Kit (Merck KGaA).

### Ab discovery and recombinant expression

Human mAbs were isolated from plasma cells or memory B cells of SARS-CoV or SARS-CoV-2 immune donors, as previously reported. Recombinant antibodies were expressed in ExpiCHO cells at 37°C and 8% CO2. Cells were transfected using ExpiFectamine. Transfected cells were supplemented 1 day after transfection with ExpiCHO Feed and ExpiFectamine CHO Enhancer. Cell culture supernatant was collected eight days after transfection and filtered through a 0.2 µm filter. Recombinant antibodies were affinity purified on an ÄKTA xpress FPLC device using 5 mL HiTrap™ MabSelect™ PrismA columns followed by buffer exchange to Histidine buffer (20 mM Histidine, 8% sucrose, pH 6) using HiPrep 26/10 desalting columns.

### MAbs pseudovirus neutralization assay

MLV-based SARS-CoV-2 S-glycoprotein-pseudotyped viruses were prepared as previously described (Pinto et al., 2020). HEK293T/17cells were cotransfected with a WT, B.1.1.7 or ΔH69/V70 SARS-CoV-2 spike glycoprotein-encoding-plasmid, an MLV Gag-Pol packaging construct and the MLV transfer vector encoding a luciferase reporter using X-tremeGENE HP transfection reagent (Roche) according to the manufacturer’s instructions. Cells were cultured for 72 h at 37°C with 5% CO_2_ before harvesting the supernatant. VeroE6 stably expressing human TMPRSS2 were cultured in Dulbecco’s Modified Eagle’s Medium (DMEM) containing 10% fetal bovine serum (FBS), 1% penicillin–streptomycin (100 I.U. penicillin/mL, 100 µg/mL), 8 µg/mL puromycin and plated into 96-well plates for 16–24 h. Pseudovirus with serial dilution of mAbs was incubated for 1 h at 37°C and then added to the wells after washing 2 times with DMEM. After 2–3 h DMEM containing 20% FBS and 2% penicillin–streptomycin was added to the cells. Following 48-72 h of infection, Bio-Glo (Promega) was added to the cells and incubated in the dark for 15 min before reading luminescence with Synergy H1 microplate reader (BioTek). Measurements were done in duplicate and relative luciferase units were converted to percent neutralization and plotted with a non-linear regression model to determine IC50 values using GraphPad PRISM software (version 9.0.0).

---

**Supplementary Figure 1.**
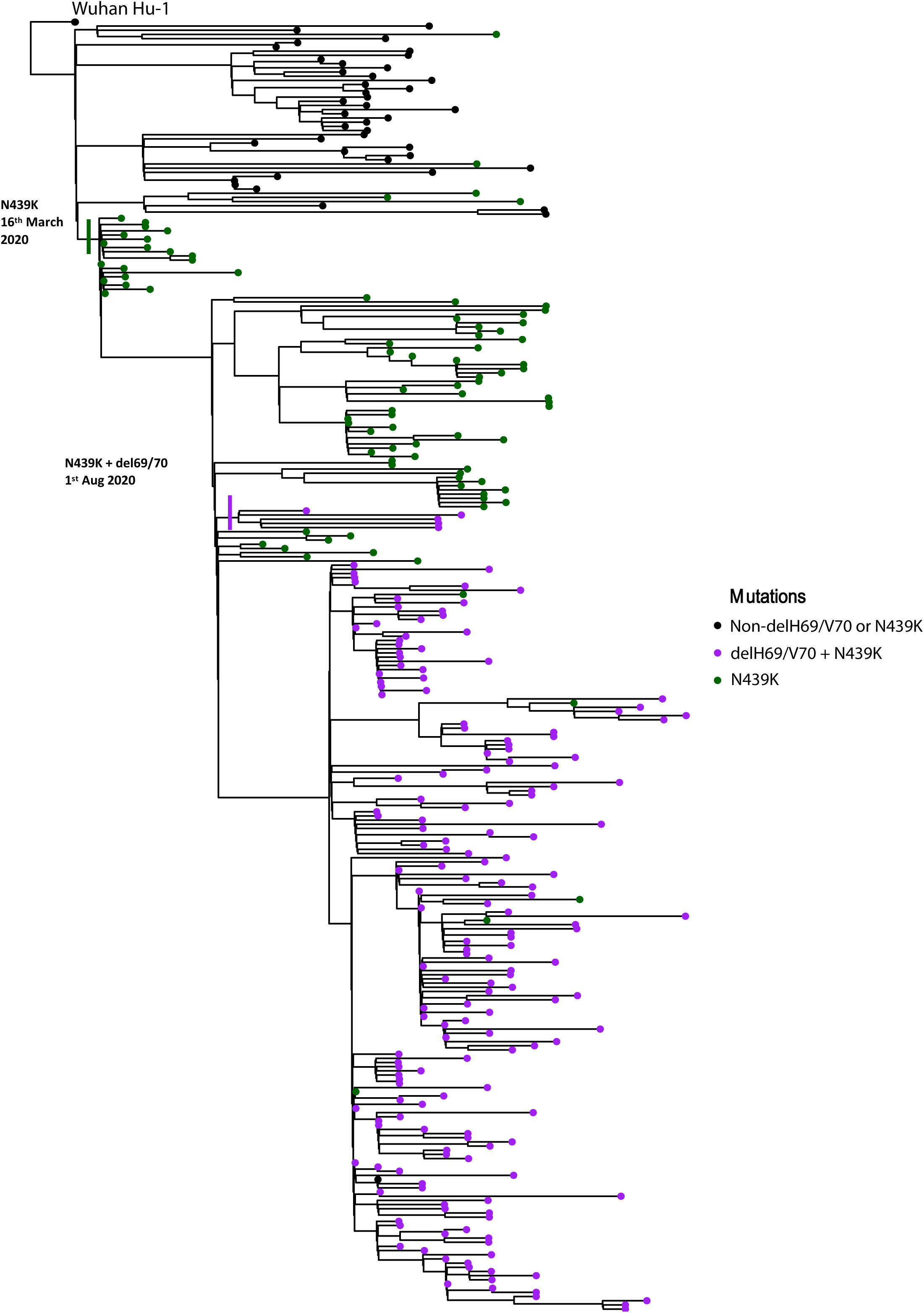
Maximum likelihood phylogeny of global sequences carrying Spike mutant N439K. All sequences in the GISAID database containing S:439K (12094 sequences, 18^th^ February 2021) were downloaded, realigned to Wuhan-Hu-1 using MAFFT and deduplicated. Viruses carrying the Spike deletion ΔH69/V70 emerged and expanded from viruses with S:439K, predominantly across the United Kingdom and Europe.

**Supplementary Figure 2:**
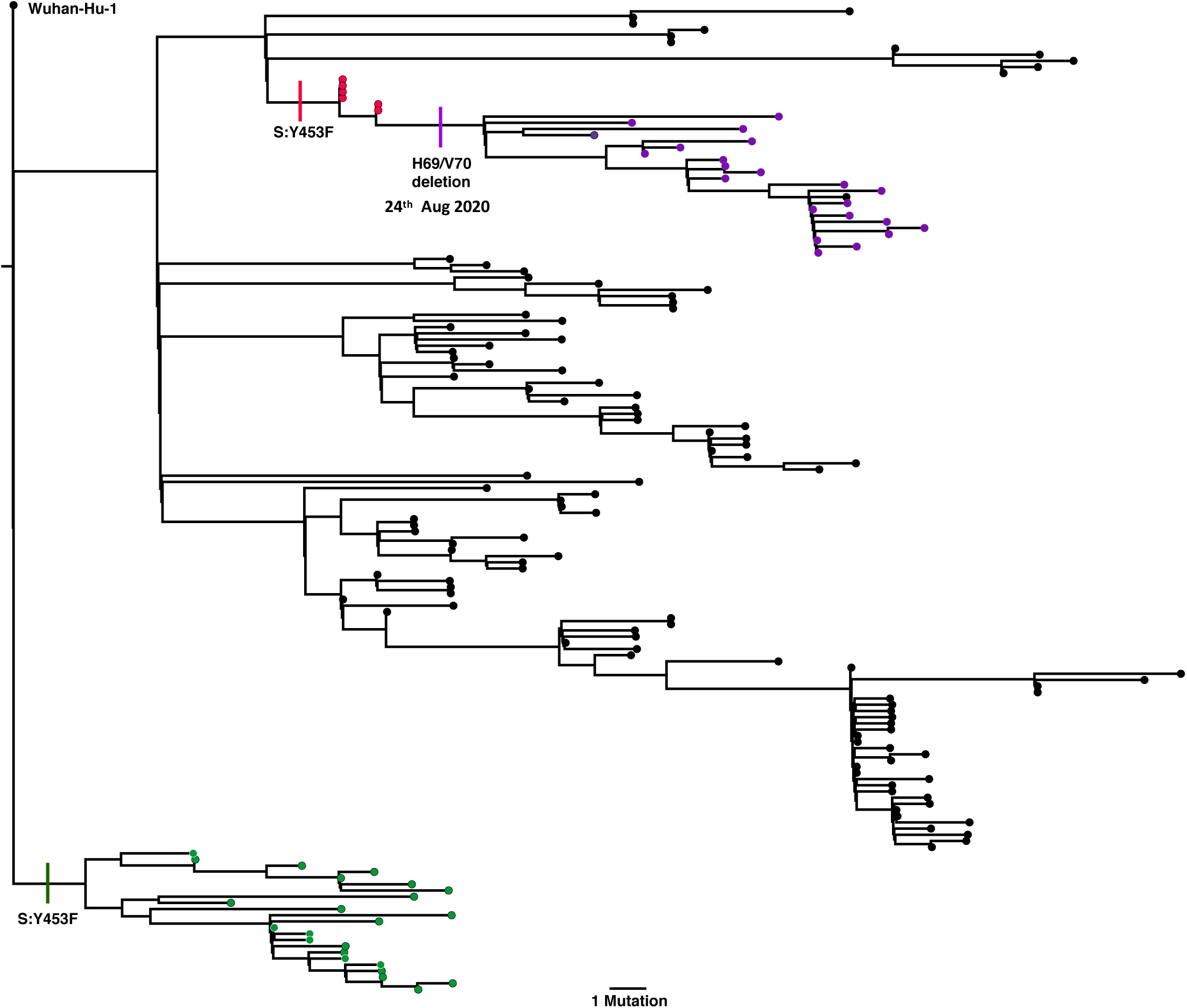
Maximum likelihood phylogeny phylogeny of SARS-CoV-2 sequences carrying Spike mutant Y453F. 753 sequences in the GISAID database (accessed 14^th^ February 2021) were downloaded and realigned to Wuhan-Hu-1 using MAFFT. Two distinct lineages carrying the mink-associated Spike Y453F mutations can be seen in Danish (red) sequences, with a separate lineage isolated only in Netherlands (green). After acquiring the Y453F mutation, Danish mink also appeared to acquire the Spike deletion ΔH69/V70 (purple).

**Supplementary Figure 3:**
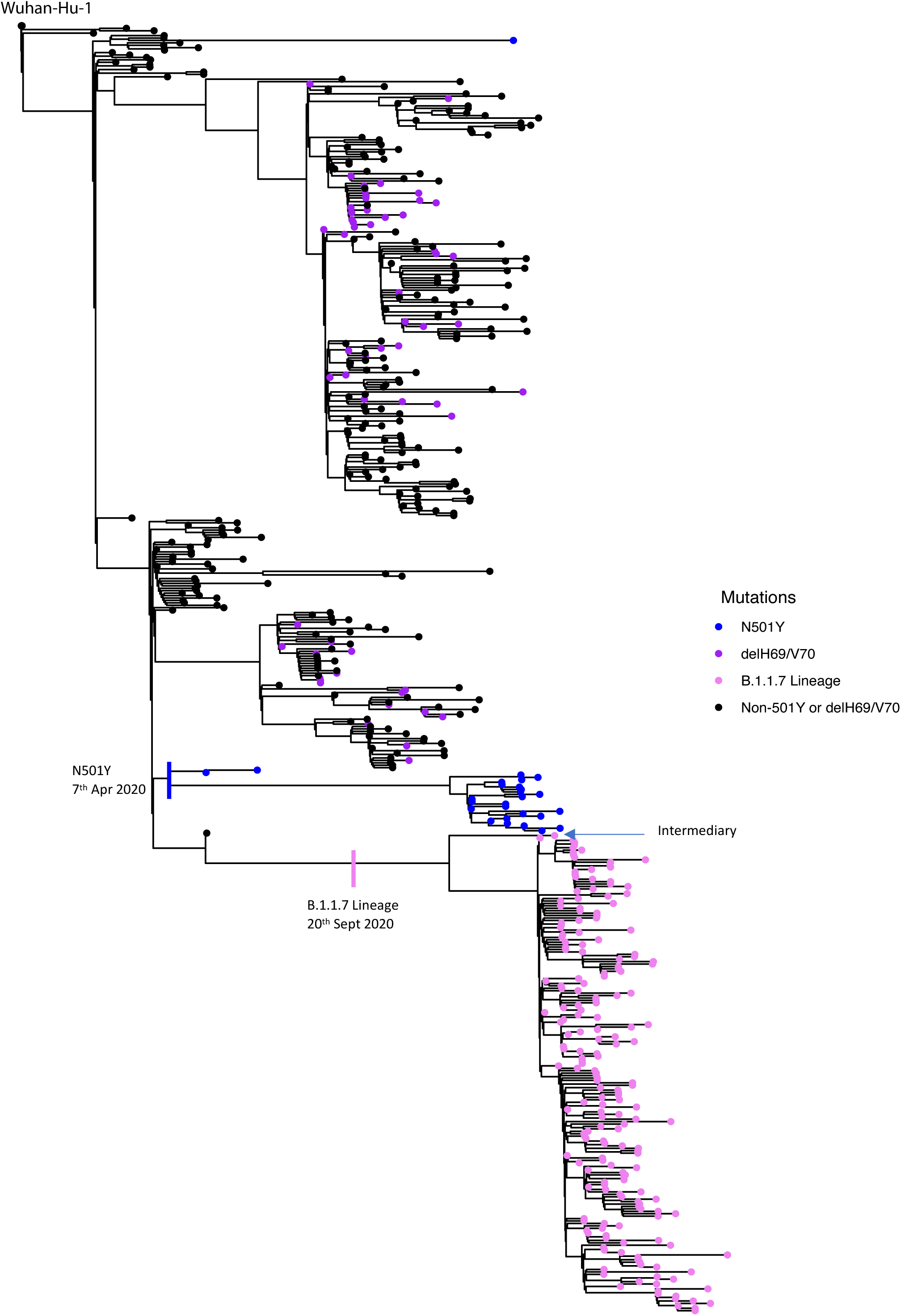
Maximum likelihood phylogeny of UK viruses bearing Δ69/70 and N501Y mutations. Two distinct lineages of the ΔH69/V70 were observed to expand in the UK, separately from the 501Y lineage. Prior to expansion of the B.1.1.7 lineage, clusters of infections bearing either N501Y or ΔH69/V70 were observed. Alongside expansion of the B.1.1.7 lineage, is a population in Wales that carries 501Y, but no ΔH69/V70. An intermediary was detected alongside the B.1.1.7 lineage (indicated on phylogeny) which had only a subset of the mutations that make up B.1.1.7 (ΔH69/V70, N501Y, A570D and D1118H).

**Supplementary Figure 4:**
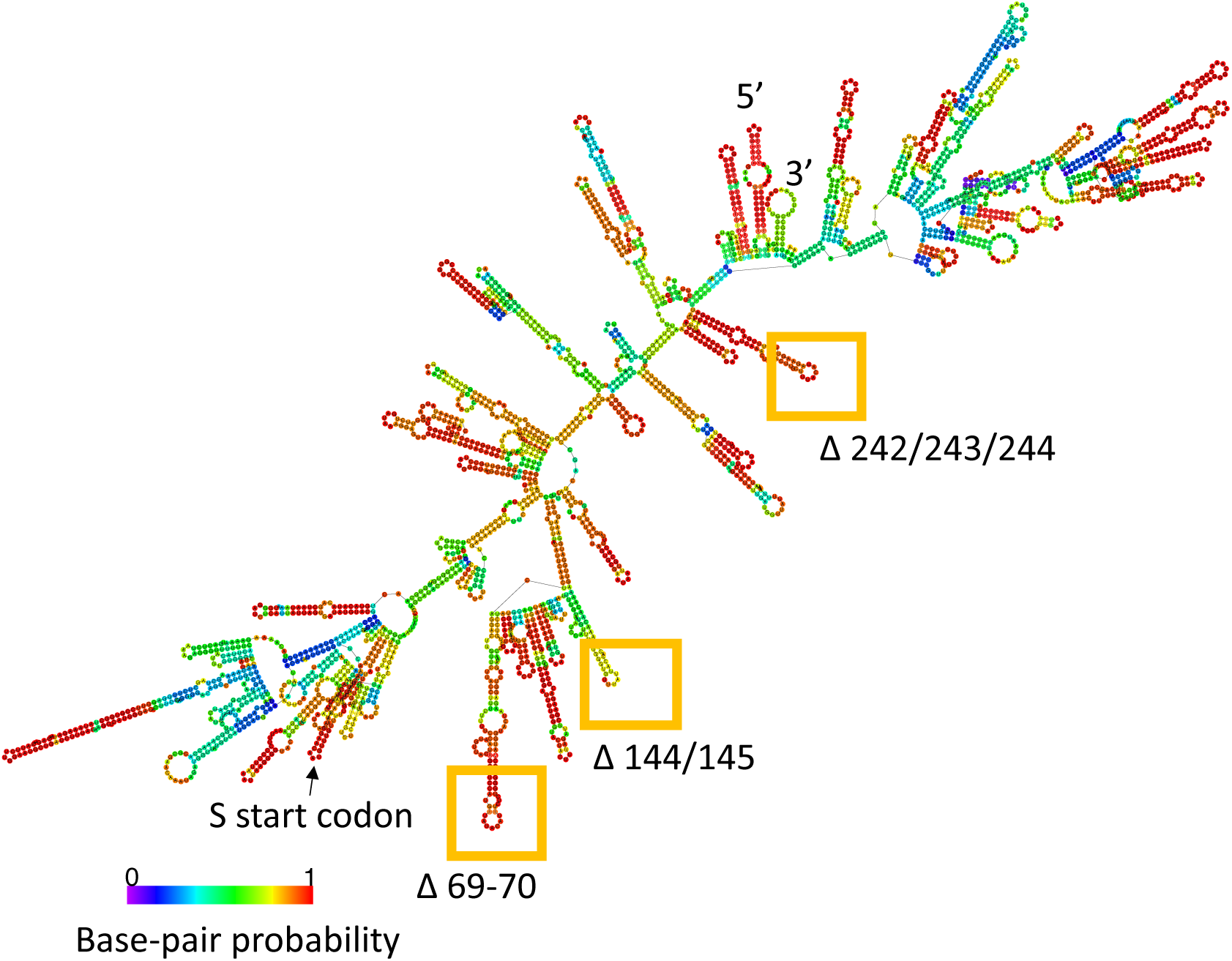
The positions of common deletion mutations on the RNA structure of the Spike Δ69-70 region of the gRNA. The optimal secondary structure was generated from a consensus alignment of human SARS-CoV2 RNAs using RNAalifold. Figure shows nucleotides 20277-23265. Base-pair probability, representative of the breadth of the structural ensemble that could be adopted by the RNA, is shown in colour according to the key.

**Supplementary Figure 5:**
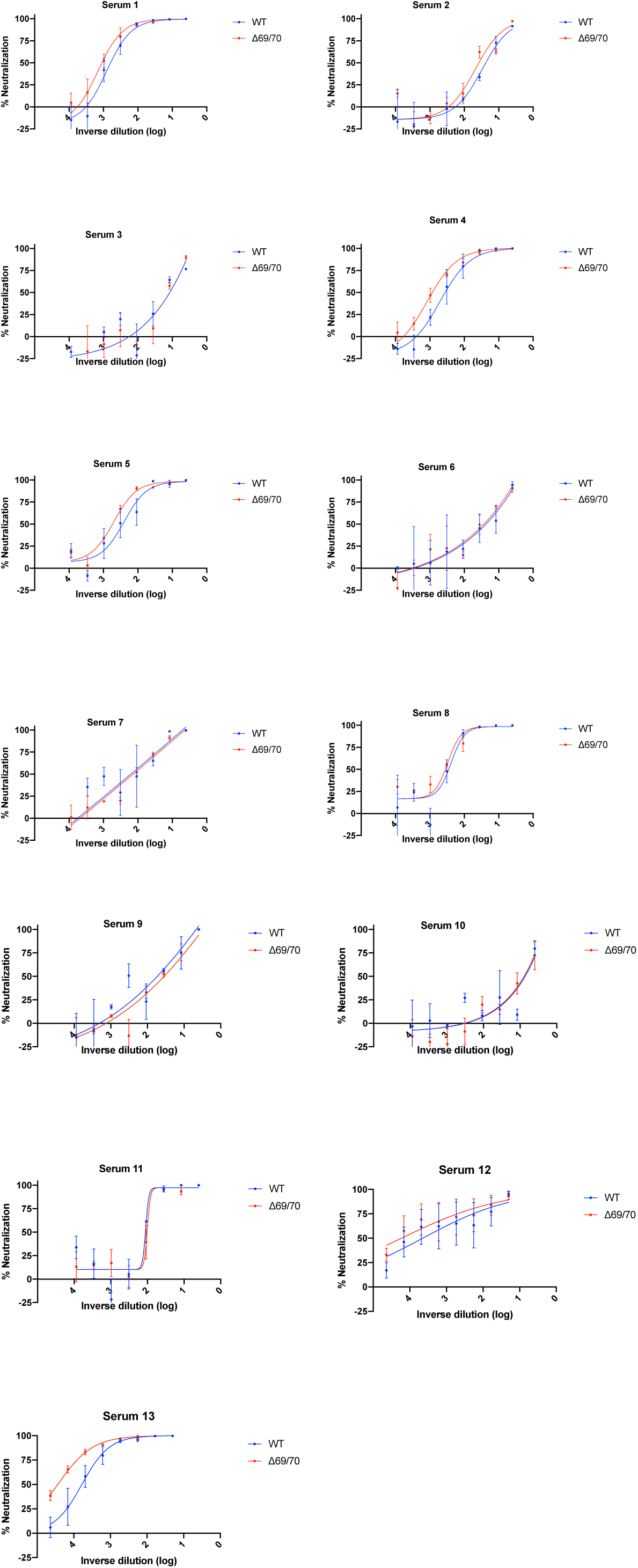
Neutralization of pseudovirus virus bearing Spike ΔH69/V70 and wild type (all In Spike D614G background) by convalescent sera from 13 donors who showed neutralisation. Indicated is serum log_10_ inverse dilution against % neutralisation. Where a curve is shifted to the right this indicates the virus is less sensitive to the neutralising antibodies in the serum. Two of fifteen original sera were non neutralising. Data points represent means of technical replicates and error bars represent standard deviation.

**Supplementary Figure 6:**
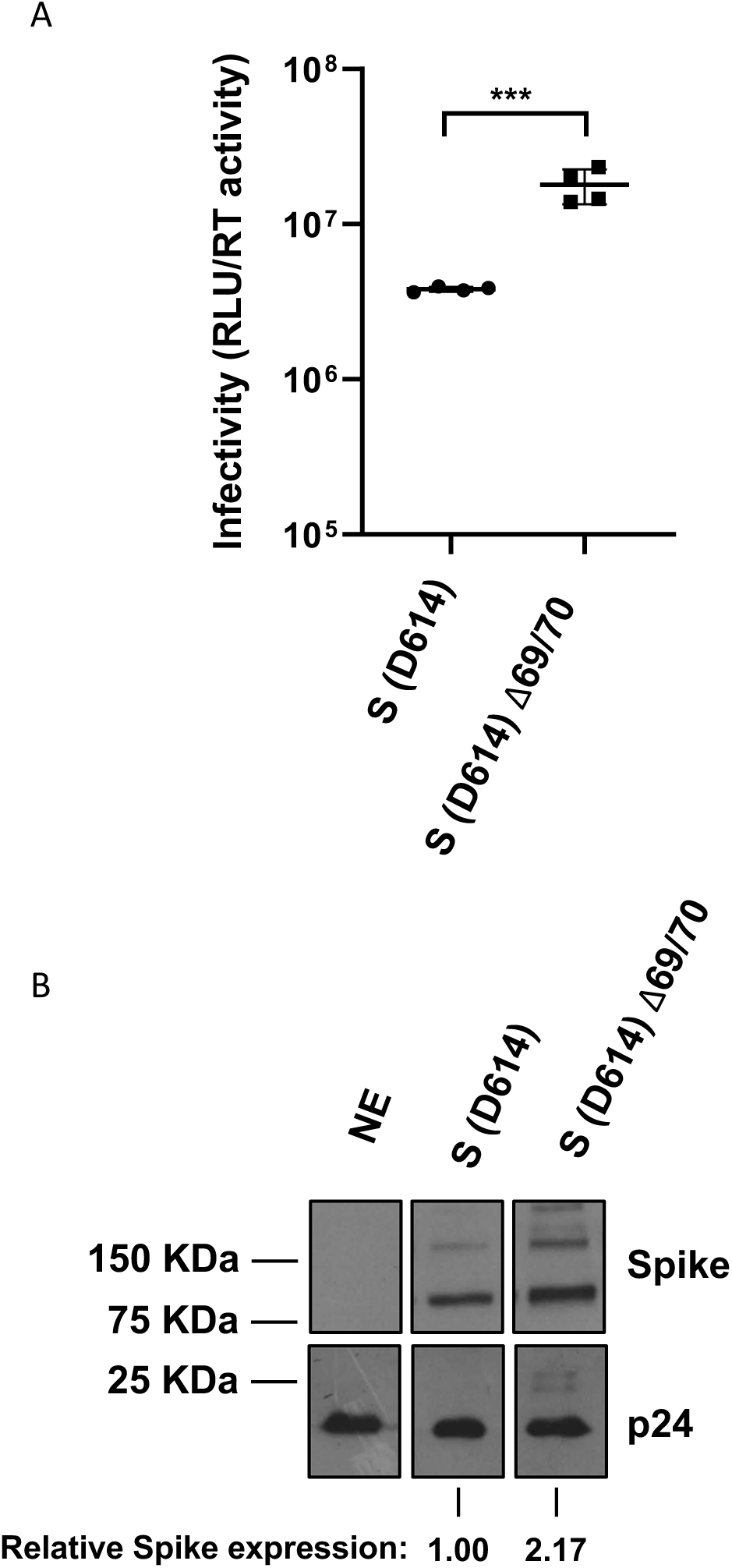
Infectivity of spike ΔH69/V70 in a background of D614 (Wuhan). Purified pseudotypes, as indicated, were used to infect human ACE2 expressing HEK293 cells, with luciferase readings read at 72 hours post infection. Experiments were performed in biological quadruplicate with the mean and standard deviation plotted. Results are representative of experiments performed two times. Statistical significance was assessed using an unpaired t-test (ns; non-significant, ***; <0.005). Western blot of purified pseudotype virus. Spike and HIV pseudotype abundances were assessed using Flag and p24 antibodies, respectively. Relative spike expression was calculated by densitometry using Image J. Briefly, inverted pixel intensities for spike and p24 bands were first normalised to a background region of the gel. Spike protein intensities were then normalised to p24 intensity before mutant protein expression was calculated as a factor of wild-type protein. NE: no envelope/spike

**Supplementary Figure 7:**
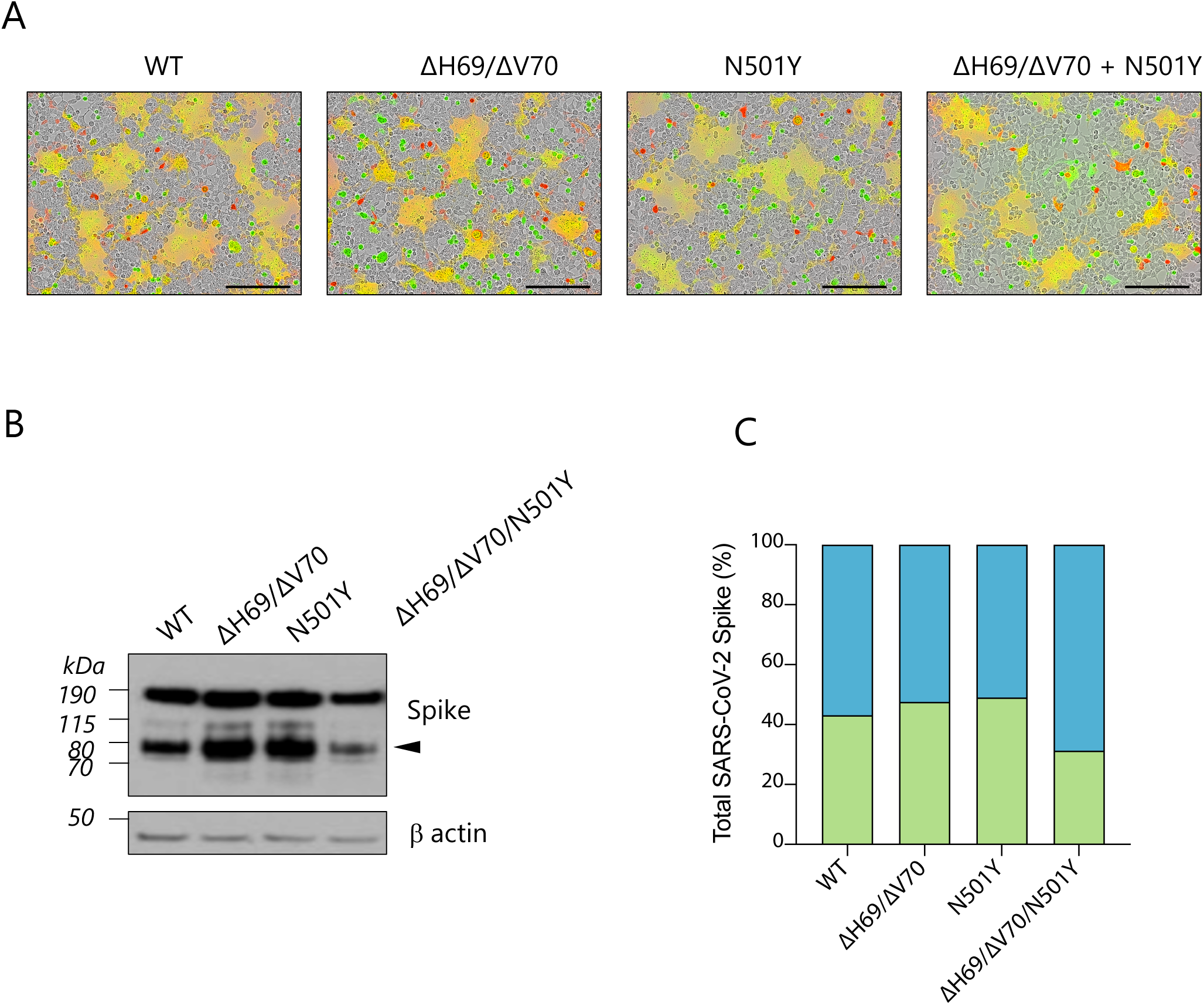
Cell Fusogenicity in the presence of Spike mutants ΔH69/V70 and N501Y. **A.** Reconstructed images at 12 hours of 293T cells co-transfected with the indicated Spike mutants and mCherry expressing plasmid mixed with green dye-labelled Vero cells. Scale bars represent 200 mm. Green colour identifies the acceptor cells while red colour marks donor cells. Merged green-red colours indicate the syncytia. **C.** Representative western blot of cells transfected with the indicated Spike mutants (detected with anti-Spike antibody). The cleaved Spike identifies the S2 subunit and is indicated with the arrowhead. b actin is shown as loading control. **D.** Quantification analysis of cleaved (green bars) and uncleaved (blue bars) Spike shown in (C) normalised by b actin.

**Supplementary Figure 8:**
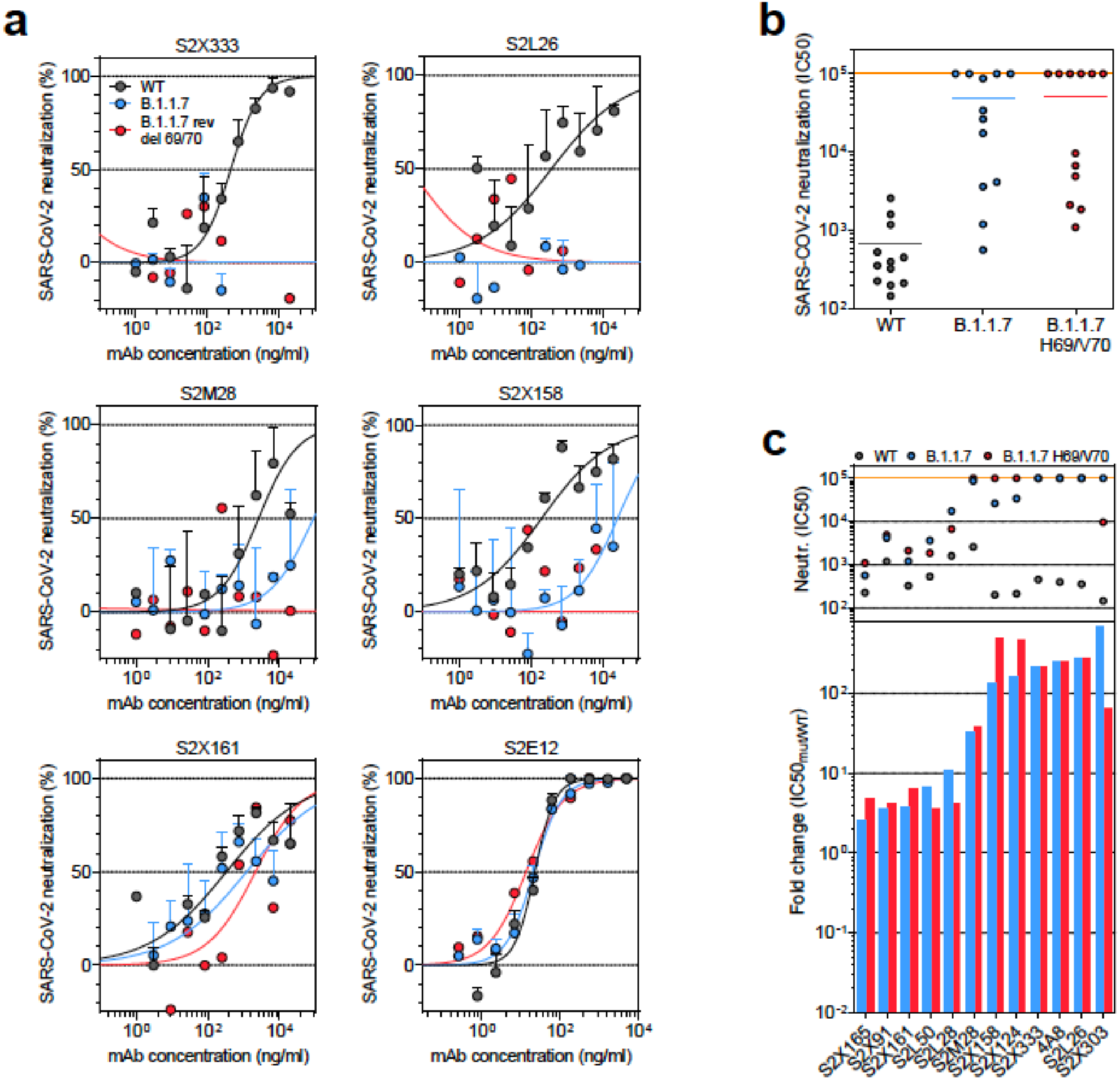
Neutralisation and binding by a panel of NTD-specific mAbs against WT, B.1.1.7 and B.1.1.7 ΔH69/V70 mutant SARS-CoV-2 viruses. **A.** Neutralisation of WT (black), B.1.1.7 (blue) and B.1.1.7 ΔH69/V70 mutant (red) pseudotyped SARS-CoV-2-MLVs by 6 selected mAbs from one experiment. **B.** Neutralisation of WT, B.1.1.7 and B.1.1.7 ΔH69/V70 SARS-CoV-2-MLVs by 13 mAbs targeting NTD. Shown are the mean IC50 values (ng/ml) from one experiment. The higher the IC50 the less sensitive the virus to antibodies. **C.** Neutralisation shown as mean IC50 values (upper panel) and mean fold change of B.1.1.7 (blue) or B.1.1.7 ΔH69/V70 (red) relative to WT (lower panel) of the 13 NTD mAbs tested. Lower panel shows IC50 values from one experiment.

**Supplementary Figure 9:**
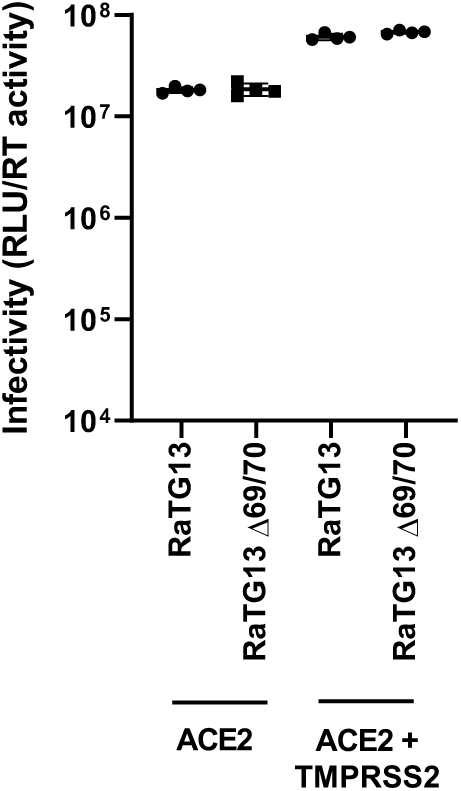
Deletion of H69V70 in the bat sarbecovirus RaTG13 spike protein does not increase infectivity in cells overexpressing ACE2 or TMPRSS2. Single round infection by luciferase expressing lentivirus pseudotyped with RaTG13 Spike protein on 293T cells transduced with ACE2 or ACE2 and TMPRSS2. Experiments were performed in biological quadruplicate with the mean and standard deviation plotted. Results are a single experiment. tNE: no envelope.

